# Anthropocene genetic diversity loss in the marine tropics

**DOI:** 10.1101/2025.09.11.675623

**Authors:** René D. Clark, Brendan N. Reid, Eric Garcia, Marial Malabag, Robin S. Waples, Rene A. Abesamis, Jemelyn Grace P. Baldisimo, Abner A. Bucol, Kyra S. Fitz, Sharon F. Magnuson, Richard N. Muallil, Cleto L. Nanola, Roy Roberts, John C. Whalen, Christopher E. Bird, Kent E. Carpenter, Malin L. Pinsky

## Abstract

Genetic diversity is a crucial component of biodiversity, and as such, its maintenance and preservation is of high conservation concern. Tropical environments are undergoing intense rates of environmental change, and these changes may be driving large declines in genetic diversity. However, data on genetic diversity are highly skewed towards temperate regions. The degree to which diversity loss has occurred in tropical species, particularly marine species, remains an open and important question. Here, we directly compare genomic data from modern and museum collections of two commercially-harvested nearshore marine fishes (*Equulites laterofenestra* and *Gazza minuta*) gathered from a single location in the Philippines, spanning a century of intense environmental change. These data reveal a marked loss in genetic diversity and evidence for multiple orders of magnitude reductions in effective population size (*N*_*e*_) in both species, indicating substantial genomic erosion. Such a decline highlights the long-lasting genomic consequences of anthropogenic activity and sheds light on an, until-now, invisible loss of diversity from the most biodiverse ocean region.

**Significance Statement:** Recent reports have shown that genetic diversity in marine fishes has declined either slightly or not at all during the Anthropocene. However, almost all studies investigating marine genetic diversity loss have been from temperate latitudes, whereas tropical marine environments experience some of the most intense human impacts. Here, we directly compare genomic data from modern and historical specimens of two commercially-harvested fishes from the Philippines. We show that marine species in the tropics have already lost substantial genetic diversity and may have undergone severe bottlenecks over the past century. These results shine a light on the previously invisible loss of genetic diversity in the most biodiverse region of the ocean, and they emphasize the evolutionary consequences of the Anthropocene.

## Introduction

Genetic diversity provides the foundation for all life on Earth, underpinning speciation events, population divergence, and adaptation to novel conditions. Recently, it has been classified as an Essential Biodiversity Variable^1^, with its preservation identified as both integral to the health of the world’s oceans^2^ and a key target for global conservation goals. However, despite this well-documented value, global genetic diversity has also experienced unprecedented loss during the Anthropocene, with an estimated 6% to 16% loss of genetic diversity based on estimates from a few dozen species^3-5^.

Despite these observations, the tropics stand out as a particular and repeated gap in knowledge in the global understanding of genetic diversity and its potential loss^3,6^. Studies investigating temporal genetic diversity loss are highly geographically and taxonomically biased, with research to date tracking evolutionary change in fishes, birds, and mammals predominantly from North America, Europe, and other temperate regions^3,5,7^. However, it is tropical marine environments that experience some of the most intense human impacts and biodiversity risks^8^.

Limited dispersal, faster generational turnover, and more fragmented population structure in the tropics^9,10^ may also result in tropical taxa being more susceptible to bottlenecks and loss of genetic diversity due to human-induced change. These factors may counteract the larger effective population sizes (*N*_*e*_), weaker genetic drift, and high gene flow experienced by many marine species that otherwise make genetic loss both unlikely and difficult to detect with sufficient power^11-13^.

Empirical evidence for genetic diversity loss in marine fishes at all latitudes remains mixed, however, partly due to the low power of existing studies^14-16^. Temporal genomic studies that can directly observe evolutionary change offer a uniquely powerful approach to investigating genetic diversity loss because historical specimens provide a window to the past and enable studies to establish genomic baselines from a time period before accelerated anthropogenic change^7^. The extent to which anthropogenic impacts can translate into substantial, long-lasting genomic consequences in the wild, and the conditions under which such change is most likely to take place, remain important questions. Furthermore, as tropical marine habitats are some of the most biodiverse and threatened ecosystems on the planet^17^, a better understanding of their unique evolutionary dynamics is important to effectively safeguard a biologically diverse future.

Here we leveraged a unique and extensive museum collection to provide one of the first temporal estimates of genetic diversity loss in tropical marine fishes and assess the vulnerability of tropical marine fish to anthropogenic change. Museum collections provide an invaluable glimpse into the past, and they are also emblematic of pervasive global scientific disparities. Natural history museums are disproportionately found in the U.S. and Western Europe, and the origins of their collections are often intertwined with colonialism^18^. The historical samples utilized here are currently curated by the Smithsonian Institution. They came from an early 20th-century research expedition by the U.S. Research Vessel *Albatross* to the Indo-Pacific, shortly after the United States forcefully acquired the Philippines as a territory at the end of the Spanish- and Philippine-American Wars^19,20^. We sequenced individuals from two ponyfish species, *Equulites laterofenestra*^21^ and *Gazza minuta*^22^ that were originally collected near Hamilo Cove in

Batangas, Philippines in 1908 (Fig. 1). Both species inhabit sandy nearshore coastal waters, share similarly short generation times of approximately 1.4 years, and are frequently targeted by fishers^23,24^. While the Hamilo coastline remains relatively intact due to its proximity to Mount Palay-Palay National Park and restricted military zones, populations in this location are still likely to have experienced intense environmental change over the past century. Nasugbu, directly south of Hamilo Cove, is a densely populated region with a robust fishing and agriculture presence^25^. Hamilo Cove is also situated near the mouth of Manila Bay, a body of water in which sustained industrialization throughout the 20th and 21st centuries has driven soil erosion, industrial waste discharge, and the deterioration of both water and habitat quality^26,27^. Additionally, agricultural wastes from fish cages in nearby Laguna de Bay contribute to nutrient loading and water pollution, further exacerbating ecological degradation^28^.

**Figure 1.**
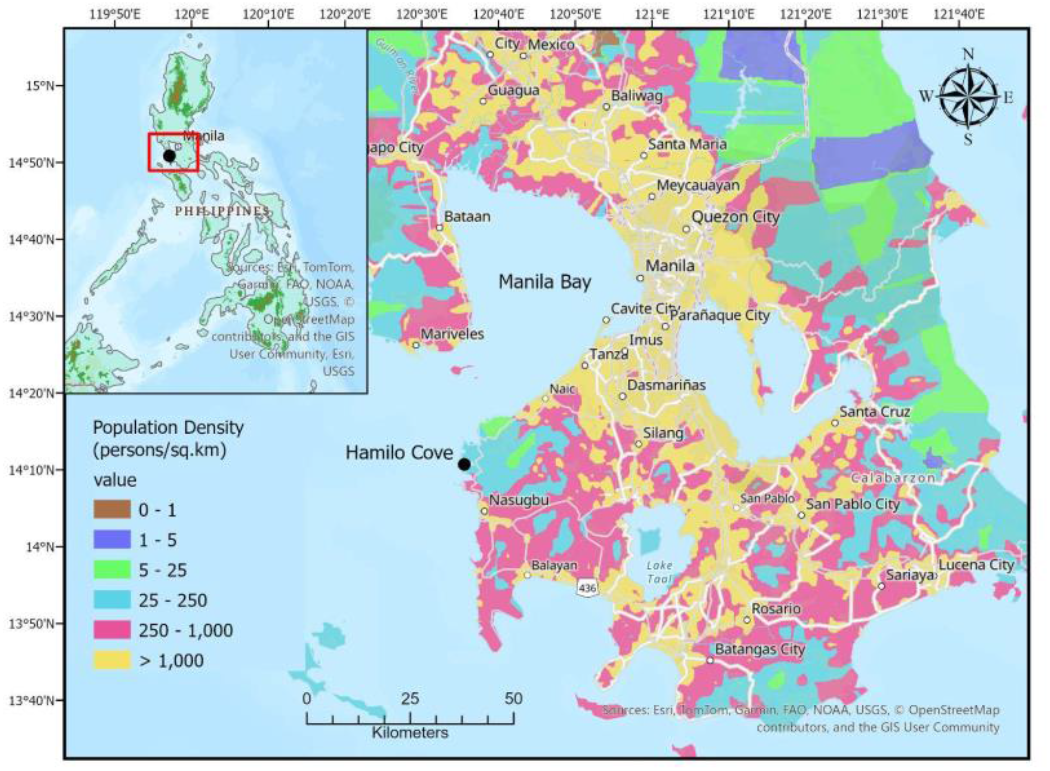
Map of the sampling location in the Philippines (Hamilo Cove) and the surrounding area. Inset map depicts the Philippines with the highlighted red box expanded for detail. Color-coding corresponds to population density (persons/km^2^), with data from the Gridded Population of the World collection (GPWv4).

We obtained capture-enriched DNA sequencing data from historical specimens from *E. laterofenestra* and *G. minuta* collected in 1908 and paired these with modern samples collected from the same location in 2018 (Fig. 1). Together, these samples allowed us to investigate the recent demographic history of these species by enabling a direct comparison of *N*_*e*_ and genetic diversity through time. In total, we genotyped over 180 individuals (29 historical, 155 contemporary) and found substantial genetic diversity loss and evidence for recent population bottlenecks, suggesting the past century of environmental decline had long-lasting evolutionary consequences for these and similar marine tropical fish populations.

## Results

We combined hybridization capture^29^ and Illumina sequencing to effectively and efficiently generate genomic data for all historical and contemporary individuals. All reads were mapped to species-specific *de novo* reference assemblies. We obtained an average read depth of 77.6 across called single nucleotide polymorphisms (SNPs) in the historical samples and an average read depth of 82.3 in the contemporary samples (Table S1-S2). *G. minuta* individuals were genotyped at 1,149,563 sites (9,641 were retained after filtering) while *E. laterofenestra* individuals were genotyped at 7,119,339 sites (53,625 post-filtering). We filtered out exogenous DNA and identified and removed individuals putatively belonging to cryptic species (SI Appendix, Fig. S1) as well as those with anomalously high heterozygosity^30^, leaving us with a total of 184 individuals (*G. minuta*: 18 historical, 63 contemporary; *E. laterofenestra*: 11 historical, 92 contemporary). Further details on sampling protocols, library preparation methods, and sequencing can be found in the Materials & Methods (SI Appendix, Fig. S2 and S3, Tables S1 and S2).

Overall, our results indicated that the same populations had been sampled through time. Principal components analysis (PCA) and ADMIXTURE^31^ revealed limited change in genomic composition for both species through time (Fig. 2, SI Appendix, Fig. S4). Contemporary and historical individuals clustered together along PCs 1 & 2 and were assigned to the same genomic population in ADMIXTURE, where K = 1 was best supported. Temporal F_ST_ was also relatively low (*G. minuta*: 0.0047, 95% CI = 0.0033, 0.0062; *E. laterofenestra*: 0.0065, 95% CI = 0.0058, 0.0072).

**Figure 2.**
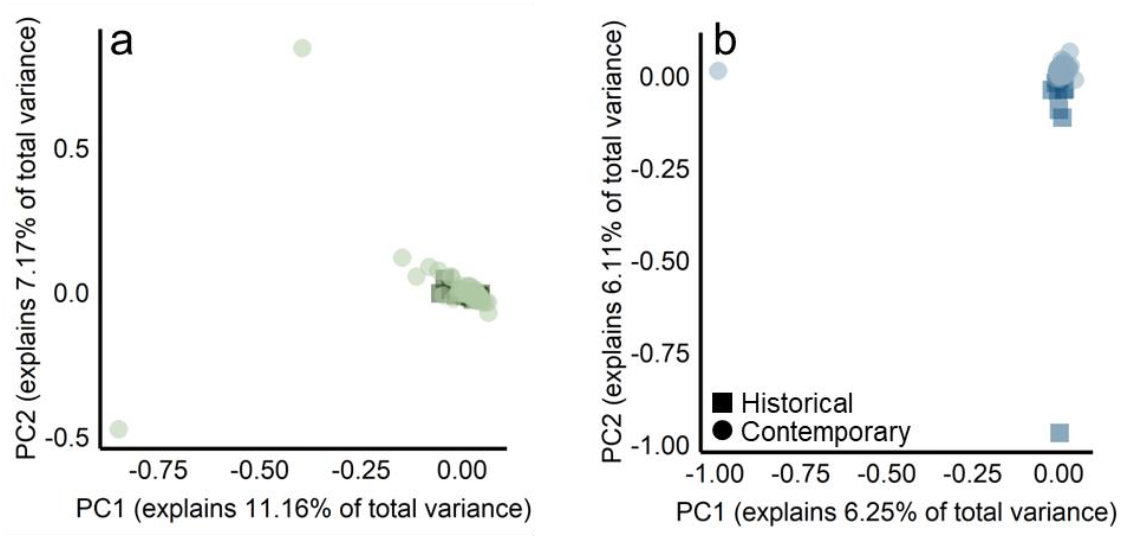
PCAs of historical and contemporary individuals for (a) *G. minuta* and (b) *E. laterofenestra*. Historical individuals are represented by dark squares; contemporary individuals are represented with lighter circles.

In both species, genetic diversity declined substantially over time (Fig. 3). On average, pairwise diversity (π) declined 4.3% (range = 3.4-5.1%), observed heterozygosity (*H*_*O*_) declined 4.3% (range = 2.7-5.8%), and expected heterozygosity (*H*_*E*_) declined 5.9% (range = 5.8-6.0%). These losses were significantly greater than expected for a null model of no genomic difference between time points for *E. laterofenestra* (SI Appendix, Fig. S5). Two-dimensional site frequency spectra (2D-SFS) for each species also revealed considerable temporal change, with many mid-frequency alleles in the historical time points reduced to low frequency in the modern day, particularly in *G. minuta* (SI Appendix, Fig. S6). Furthermore, *E. laterofenestra* (but not *G. minuta*) also displayed a small positive shift in genome-wide Tajima’s D through time, a classic signature of recent population contractions (SI Appendix, Fig. S7). Finally, the inbreeding coefficients (F_IS_) in *G. minuta* declined slightly, albeit not significantly, over time, which may indicate heterozygote excess and reduced *N*_*e*_ in the contemporary population^32^ (Fig. 3).

**Figure 3.**
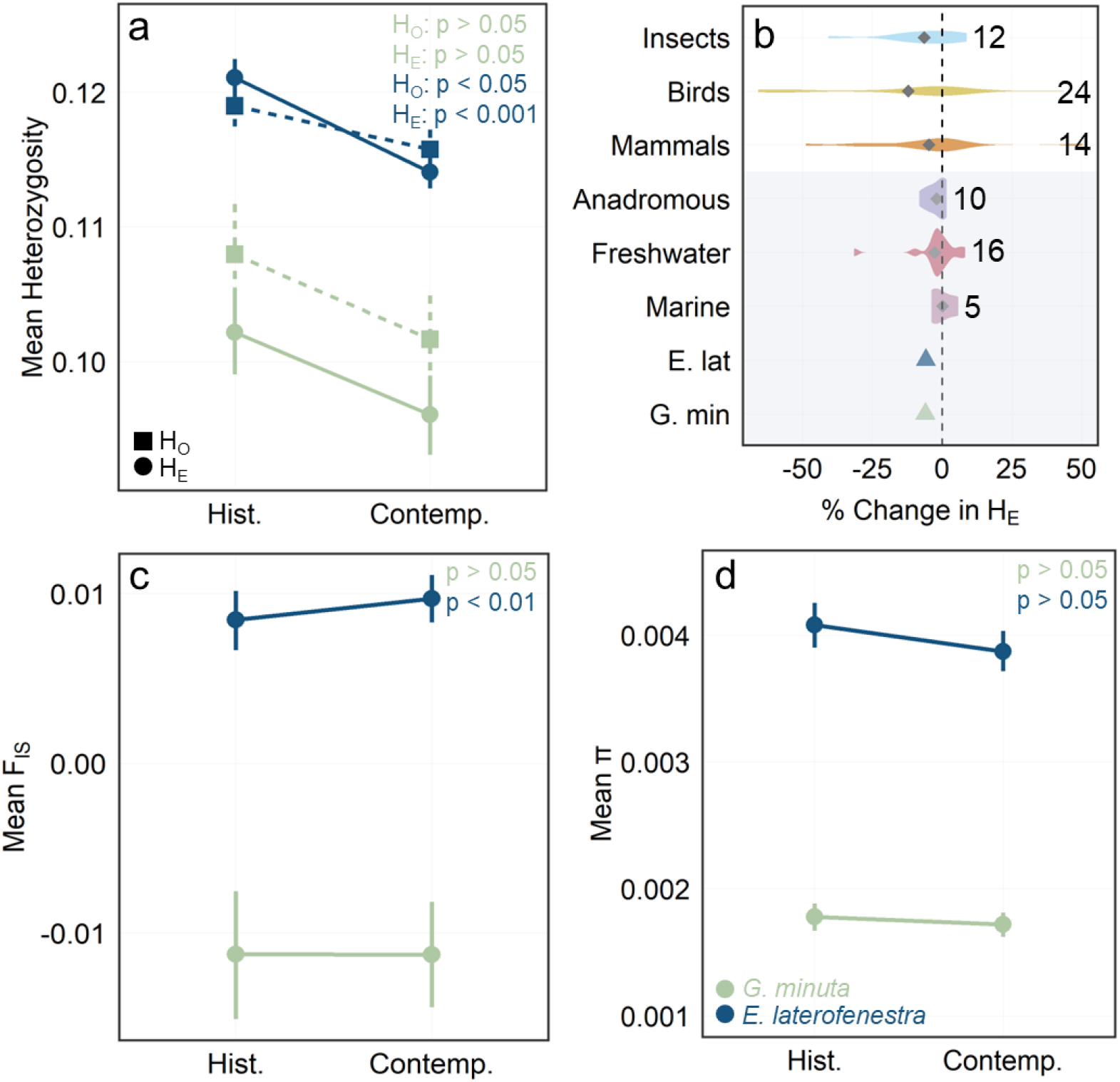
Genetic diversity estimates for each species and both historical and contemporary time points (green: *G. minuta*, blue: *E. laterofenestra*): (a) *H*_*E &*_ *H*_*O*_, (b) % change in *H*_*E*_, (c) π, (d) F_IS_. Error bars represent 95% CIs. For each diversity estimate, reported empirical p-values indicate the significance of the observed change in diversity compared to a null model of no genetic change through time. Panel b depicts the mean % change in *H*_*E*_ over time for *G. minuta, E. laterofenestra*, and a range of other species (data from Leigh et al. 2019). The gray box surrounds fish taxa. Positive values indicate an increase in *H*_*E*_ over time, while negative values indicate a decrease. The dashed vertical line at zero indicates no change over time. Numbers next to each violin plot represent sample sizes. Gray diamonds represent mean % change in *H*_*E*_ for each taxonomic group.

Strikingly, these declines in tropical marine genetic diversity are larger than any observed in temperate marine fish populations, which have maintained diversity through time (Fig. 3b)^3,5^. Instead, these tropical declines are commensurate with the losses documented for terrestrial species. This tropical loss of diversity was unanticipated due to the higher *N*_*e*_s and weaker drift generally expected by marine populations^12^. As larger populations are less sensitive to drift-induced genetic diversity loss than smaller populations^33^, the scale of the diversity loss observed here suggests that the population declines in these tropical fishes were substantially more severe than in terrestrial birds and mammals.

To identify signatures of selection through time, we screened SNP genotypes for temporal outliers using BayeScan^34^. Only one temporal outlier was found, and it appeared in only one species, *G. minuta* (SI Appendix, Fig. S8). While our reduced representation sequencing design hindered our ability to detect selection because it only genotyped a fraction of each species’ genome, these results tentatively suggest that the large and repeated temporal declines in genetic diversity were not complemented by strong signatures of selection.

The losses in genetic diversity we observed suggest that these species have undergone pronounced genomic bottlenecks over the past century of substantial human impacts. We therefore estimated recent demographic history with momi2^35^ by fitting models of changes in *N*_*e*_ against the site frequency spectrum (SFS) for each species and time point. momi2 is particularly adept at detecting recent change in *N*_*e*_, especially when detailed linkage information does not exist, as is the case here^36^. We tested among fifteen demographic models: (1) no temporal change in *N*_*e*_, (2) a recent change in *N*_*e*_ between the historical and contemporary collection time points, (3) a historical change in *N*_*e*_ within 400 years prior to the historical collection time point, (4) an “ancient” change in *N*_*e*_ 10,000 - 1,000,000 years ago, (5) a recent and “ancient” change in *N*_*e*_, (6) a historical and “ancient” change in *N*_*e*_, (7) a recent and historical change in *N*_*e*_, (8) all three changes in *N*_*e*_, and (9-15) each of the previous seven scenarios as gradual (exponential) rather than abrupt (instantaneous) changes in *N*_*e*_ (SI Appendix, Fig. S9). Models were structured in this manner to specifically test for a recent bottleneck (between the collection time points) while also accounting for the possible presence of ancestral changes in *N*_*e*_ during the Pleistocene, which have been well-documented in tropical marine taxa^37^. For both species, when including only contemporary data in the model, the model with both a recent and “ancient” gradual change in *N*_*e*_ performed best (Fig. 4, Table 1, SI Appendix, Tables S3 and S4). In both cases, *G. minuta* and *E. laterofenestra* were predicted to have experienced a deeper-time population expansion, followed by a more recent, dramatic bottleneck and substantial decline in *N*_*e*_ within the last century (Table 1). When including historical data, models that included both a historical and “ancient” change performed best for both species (SI Appendix, Fig. S10, Tables S5-S10), suggesting that the bottleneck may have begun before 1908. However, there was also some evidence for a recent expansion to varying degrees in both species, which may indicate difficulties resolving recent dynamics from genomic data^38^.

**Table 1.** *N*_*e*_ estimates for both species, generated from demographic modeling with momi2. The first 5 columns contain results from the most parsimonious model fit to the contemporary SFS. Estimates of ancient *N*_*e*_, historical *N*_*e*_ (*N*_*e*_ of the historical population), contemporary *N*_*e*_, and the timing of the ancient and recent changes in *N*_*e*_ (in years before present) are provided. The last 2 columns contain estimates of temporal *N*_*e*_. 95% CIs are provided in brackets.

| | Ancient $N_e$ | Historical $N_e$ | Contemp.<br>$N_e$ | Time of<br>Ancient $N_e$<br>Change | Time of<br>Recent $N_e$<br>Change | Allele<br>Frequency<br>Temporal $N_e$ | $H_E$ Loss<br>Temporal<br>$N_e$ |
| --- | --- | --- | --- | --- | --- | --- | --- |
| *G. minuta* | 63,955<br>[106 -<br>71,302] | 20,025,200<br>[1,236,161 -<br>9,964,418,183] | 60<br>[4 - 212] | 191,148<br>[163,083 -<br>996,992] | 75<br>[3 - 100] | 6,949<br>[4,402 -<br>16,492] | 639<br>[318 -<br>48,820] |
| *E. laterofenestra* | 678,668<br>[658,606 -<br>845,923] | 42,340,045<br>[12,562,286 -<br>9,981,472,336] | 10<br>[1 - 74] | 804,608<br>[10,000 -<br>999,935] | 16<br>[1 - 95] | 3,712<br>[3,222 -<br>4,378] | 662<br>[482 -<br>1,068] |

**Figure 4.**
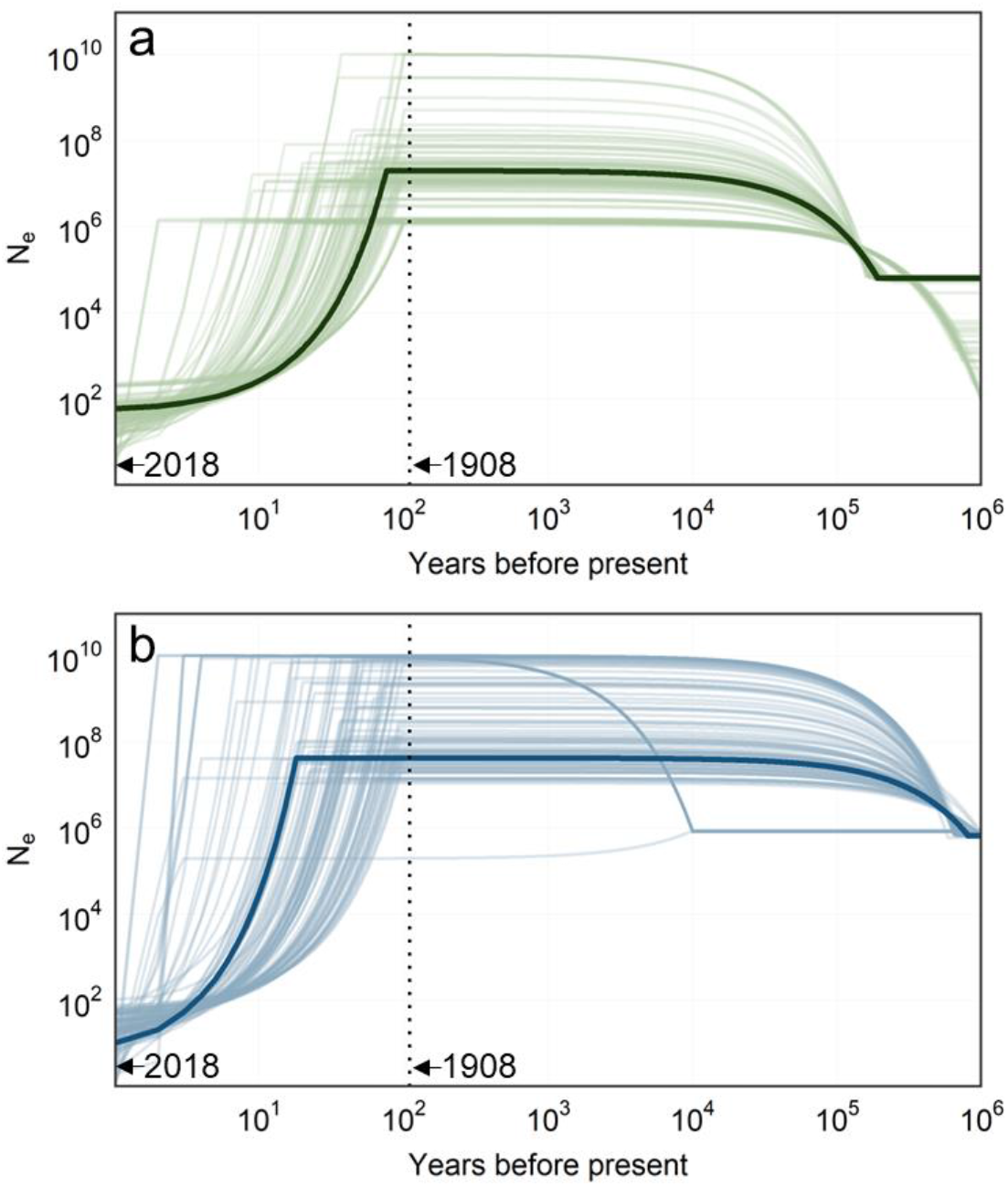
Plots of *N*_*e*_ through time for (a) *G. minuta* and (b) *E. laterofenestra*. Both axes are depicted in log scale. For each figure, the solid dark line represents estimated *N*_*e*_ from the most parsimonious model fit to the observed contemporary SF, while lighter lines represent estimated *N*_*e*_ from the most parsimonious model fit to 100 bootstrapped contemporary SFS (each line represents one bootstrap). The dashed vertical line indicates the 1908 time point. Estimates were generated using momi2; changes in *N*_*e*_ were modeled as exponential growth (or decline).

In addition to the SFS-based estimates of *N*_*e*_ provided by momi2, we calculated temporal *N*_*e*_, representing the harmonic mean *N*_*e*_ between 1908 and 2018. Temporal *N*_*e*_ was estimated two ways, using (1) the change in allele frequencies across time points^39,40^ and (2) the observed loss in heterozygosity due to drift^41^. The allele frequency change method generated temporal *N*_*e*_ estimates in the low thousands (*G. minuta*: 6,949; *E. laterofenestra*: 3,712) while the heterozygosity loss method placed temporal *N*_*e*_ at approximately 650 individuals for both species (Table 1). Harmonic means give extra weight to smaller values and are particularly sensitive to the lower *N*_*e*_s that appear after bottleneck events. Importantly, both estimators may be influenced by age and spatial structure, although the impact of age structure should be minimal given the species’ short generation times and the large number of elapsed generations^42^. Similarly, although the greater genetic structure and low-to-moderate migration rates of tropical marine species like ponyfish^43^ enhance our ability to capture local rather than global *N*_*e*_, the true local *N*_*e*_ for both populations are almost certainly smaller than our estimates, as we are likely retaining a partial signal of metapopulation *N*_*e*_.

## Discussion

Museum specimens provide unparalleled insight into past populations and can greatly enhance our ability to detect and observe evolutionary processes *in situ*. We demonstrated the power of such collections here by comparing genomic data from historical and modern samples of two tropical marine species collected from the same location 110 years apart. For both species, we found a marked loss in genetic diversity and our top demographic models indicated the occurrence of strong recent bottlenecks, suggesting the past century of intensifying anthropogenic change has resulted in substantial genomic erosion. Strikingly, the rate of diversity loss we observed was comparable to those previously identified in birds and mammals, and greater than those for temperate marine fishes^3^. This loss occurred even though marine taxa have typically been thought to have large, open populations with weak genetic drift, and thus are unlikely to experience genetic diversity decline at the same rate as smaller, more fragmented species. However, nearshore tropical reef communities tend to be less connected than temperate or pelagic taxa due to shorter pelagic larval durations and higher levels of self-recruitment^9,10,44^. Such characteristics may make tropical marine taxa more susceptible to bottlenecks and loss of genetic diversity, particularly because they limit the ability of tropical populations to replenish their genetic stock from elsewhere within the species range. Moreover, these shared features may be one reason we detected declines in genetic diversity and *N*_*e*_ across both tropical species, despite similar patterns being absent among temperate fishes.

Most temporal genomic studies on marine taxa have investigated highly abundant commercial species at temperate latitudes, often motivated by an interest in declines following periods of overfishing^7^. Commercially-fished species (e.g., cod) tend to be fished because they have high abundance, and these species therefore generally have high *N*_*e*_, weak genetic drift, and little to no evidence for genetic diversity loss^13,14,16^. Our results, in contrast, caution that such species do not represent marine taxa as a whole. Research across a wider range of non-commercial species is needed to understand the generality of diversity loss in the tropics and across all latitudes. For example, nearshore species even at temperate latitudes can display reduced connectivity and greater population structure^45^. Our findings here provide testable hypotheses and lay the groundwork for future studies in this area, as can the growing accessibility of museum data^19,46^.

Both of our species were ponyfishes (Leiognathidae) that share key traits and respond to environmental changes in similar ways. Ponyfishes often form abundant mixed-species assemblages with conserved morphologies across taxa^24^, exposing many species to the same set of environmental changes and facilitating shared evolutionary trajectories. Importantly, *G. minuta* and *E. laterofenestra* are a favored food in the Philippines and frequently caught by local artisanal fishers using gill nets or beach seines, as well as by larger commercial fishers employing bottom trawls that are more destructive to marine habitats^47^. These factors may have driven similar and large declines in abundance for both species. Additionally, ponyfish are demersal invertebrate feeders that inhabit sandy, nearshore habitats and may be more likely to directly consume anthropogenic waste runoff, further impacting population health^48^. However, because variation in food web roles, reproductive traits, and population dynamics (e.g., boom and bust cycles) may increase or decrease a species’ susceptibility to anthropogenic activity, future studies exploring temporal genomic change in other tropical marine species are needed.

Demographic modeling with momi2 suggested that, over the past millennia, both species appear to have undergone an extended historical period of population expansion followed by a recent, dramatic bottleneck (Fig. 4). The more contemporary bottlenecks likely occurred synchronously in *G. minuta* and *E. laterofenestra* and may have begun as a result of prolonged habitat fragmentation and degradation. The sampling location, Hamilo Cove, is sandwiched between regions that have been heavily degraded by humans over the past century, including densely populated towns to the north and agricultural lands to the south. Heavy deforestation of the Batangas province for agricultural (particularly livestock and sugarcane production) and residential purposes began during the Spanish colonial period (1565-1898) and continued to accelerate throughout the 20th century^49^. This loss of forest cover subsequently increased surface runoff, resulting in higher erosion and siltation rates in near-shore habitats^50^, which can negatively impact coastal ecosystems by reducing sunlight availability, decreasing primary productivity, and facilitating the restructuring of shoreline communities^51^. Hamilo Cove’s proximity to Manila and Manila Bay also heightens its ecological risk. The shoreline of Manila Bay has been heavily altered, with a marked shift from mangrove habitats to human-built coastal developments over the past century, coinciding with the timeframe of the recent population bottlenecks observed here^52^. Because of this industrial and human population growth, Manila Bay is now a “marine pollution hot spot”^53^, with high outflows of untreated sewage, industrial waste (including heavy metals), and riverine plastic pollution^54,55^ that can lead to fish kills, either directly or indirectly^56^. These environmental changes have likely contributed to the sharp reductions in *N*_*e*_ observed here, particularly as recent surveys have indicated concurrent declines in fish abundance, catch rates, and coral cover, both in Batangas and elsewhere throughout the Philippines^56,57^. Ponyfish sensitivity to these changes is consistent with the more general loss of genetic diversity in nearshore and estuarine species affected by urbanization^58^.

Although our reduced representation sequencing design was well-suited to detecting temporal changes in both genetic diversity and *N*_*e*_, such approaches often lack the power to detect signatures of selection because they examine small fractions of the genome. Future studies using low- or high-coverage whole-genome sequencing will be better suited to identifying any such signatures. We additionally dealt with small sample sizes of 11-92 individuals per time point, although one major cause of reduced sample sizes was the presence of cryptic species in our datasets. While not fully explored here, the elevated levels of marine biodiversity in the Philippines^59^, coupled with the fact that ponyfish often appear in mixed-species assemblages and can be difficult to identify externally due to shared morphological features^23^, suggest the presence of cryptic species is not surprising. Moreover, the presence, and potential loss, of cryptic species reinforces the narrative that other dimensions of diversity may also be declining in the tropics, oftentimes unnoticed without careful taxonomic identifications, and heightens the call for future work documenting complementary patterns of species loss and turnover in tropical ecosystems.

These results suggest that tropical reefs and coastal systems worldwide are experiencing ongoing genomic erosion, driven in part by over-fishing, habitat degradation, and pollution. Such declines highlight the ability of anthropogenic impacts to translate into long-lasting genomic consequences and limit future adaptive capacity, particularly in some of the most biodiverse portions of the oceans. The recognition and protection of global genetic diversity is now an international goal^1,2^, with suggestions to maintain 90% or more of genetic diversity worldwide^60^. If such targets are to be taken seriously, research investigating how, when, and to what extent genomic decline is taking place in tropical systems is urgent and necessary, before these conservation goals drift out of reach.

## Materials and Methods

### Sampling, Extraction, Library Preparation, and Sequencing

Historical fish samples for *G. minuta* and *E. laterofenestra* were obtained from the Smithsonian National Museum of Natural History (NMNH) (SI Appendix, Table S1). These samples were originally collected in 1908 by the U.S. Research Vessel *Albatross* during an expedition to the Indo-Pacific^19^. Samples from both species were collected near Hamilo Cove, Philippines. Contemporary samples were obtained from the same location in the Philippines in 2018 from public markets, landing sites, or from local fishermen (SI Appendix, Table S2). Contemporary samples were stored in 95% EtOH, while historical samples were originally maintained in high proof rum distillate before being transferred to 75% EtOH at the Smithsonian Institution.

DNA extraction took place at Old Dominion University. For each sample, DNA was extracted from 45-50 mg of muscle tissue using the Qiagen Blood & Tissue Kit following a modified protocol (see the SI Appendix, Supporting Methods for more information). Library preparation was performed by the Genomics Core Laboratory (GCL) at Texas A&M University - Corpus Christi. Capture libraries, for both historical and contemporary individuals, were prepared following Roberts^61^ with the addition of a hybridization capture protocol^29^. For additional information on both the capture bait design and library preparation protocol, see the SI

Appendix, Supporting Methods. Libraries were subjected to paired end 2 x 150 bp sequencing on an Illumina NovaSeq 6000 by Novogene Corporation Inc. (Sacramento, CA).

### Data Processing and Genotyping

Processing of sequence data was accomplished following the *pire_fq_gz_processing* GitHub repository^62^. See the SI Appendix, Supporting Methods for more details. dDocentHPC was used for read mapping, genotyping, and variant filtering. For *G. minuta*, reads were mapped to the *de novo* assembly created during the capture bait design step with BWA-mem2 v.2.2.1^63^. BWA-mem has been shown to be a particularly effective mapping algorithm for mapping historical reads that are >100 bp in length and can help reduce issues of reference or mapping bias^64-66^. For *E. laterofenestra*, reads were mapped to a *de novo* whole genome assembly created with shotgun sequencing and assembled by SPAdes v.3.15.3^67^ following the *pire_ssl_data_processing* GitHub repository^68^. Reads with a minimum mapping quality less than 30, unmapped reads, and reads with secondary or supplementary alignments were filtered with SAMtools v.1.9^69^. Historical damage patterns were quantified with mapDamage2 v.2.2.1^70^ and base quality scores were recalibrated to account for base-calling errors characteristic to historical DNA degradation (SI Appendix, Fig. S2). Variants were called with FreeBayes v.1.3.1^71^. We retained only biallelic SNPs. For information on additional filtering with VCFtools v.0.1.14^72^ and the identification and removal of cryptic species or contaminated individuals see the SI Appendix, Supporting Methods.

### Population Structure, Differentiation, and Genetic Diversity

We analyzed population structure with PCA and ADMIXTURE v.1.3^31^. For both species, the full, LD-pruned SNP dataset without cryptic species or high heterozygosity individuals was used for both PCA and ADMIXTURE. PCA was performed using PLINK v.1.9^73^. A suite of variable features drove differentiation along the PCA axes (PCs 1 & 2) for both species. For example in *Gazza minuta*, a single individual (CBat_009) with higher heterozygosity drove differentiation on PC 1, while a different high-heterozygosity individual (CBat_045) drove differentiation on PC 2. When these “outlier” individuals were removed, new highly differentiated individuals appeared along PCs 1 & 2 for both species; thus we retained the original set of individuals for PCA, ADMIXTURE, and all other downstream analyses. ADMIXTURE was run for values of K from 1-5, and the optimal value of K was identified as the one with the lowest cross-validation error. ADMIXTURE results were visualized with the *pophelper* v.2.3.1 package^74^ in R v.4.2.2^75^. As a sensitivity test, we also conducted all analyses with both the cryptic species and high heterozygosity individuals included. In addition, temporal pairwise F_ST_ estimates and 95% CIs were calculated for each species using the *hierfstat* v.0.5.11 package^76^ in R.

For both species, we assessed three measures of genetic diversity. Within each time point, we calculated observed heterozygosity (*H*_*O*_) and expected gene diversity (*H*_*E*_) with the *hierfstat* package in R. The inbreeding coefficient (F_IS_) was calculated as 1 - (*H*_*O*_/*H*_*E*_). We also calculated π using pixy v.1.2.7^77^. To distinguish between monomorphic sites and truly missing data, we calculated π using an “all sites” SNP dataset that included both monomorphic and polymorphic data and was not LD-pruned (resulting in 467,359 sites for *G. minuta*, and 485,532 sites for *E. laterofenestra*). Tajima’s D was calculated across each contig using VCFtools and the same “all sites” dataset. For all diversity estimates, we calculated 95% CIs by bootstrapping with replacement across sites 1,000x in R using the boot v.1.3.28 package^78^.

To determine the statistical significance of any observed change in diversity through time, we created null models of no genomic difference between time points. Within species, we randomly re-sampled across time points without replacement 10,000x to create permuted populations composed of a mix of historical and contemporary individuals. We then calculated the difference in *H*_*O*_, *H*_*E*_, F_IS_, and π between these permuted populations and used this distribution to derive empirical p-values.

Finally, for each species, we generated the observed 2D-SFS with easySFS.py (https://github.com/isaacovercast/easySFS) using the “all sites” dataset. To retain as many informative variants as possible, we downsampled the populations to exclude individuals with a large amount of missing data. *G. minuta* was downsampled to 14 individuals in the historical time point and 36 individuals in the contemporary time point, while *E. laterofenestra* was downsampled to 9 individuals in the historical time point and 80 in the contemporary time point. All 2D-SFS were visualized with *dadi* v.2.0.5^79^.

### Test for Selection

Within each species, SNP genotypes were screened for temporal outliers using BayeScan v.2.1^34^ with prior odds for the neutral model equal to 100. BayeScan tests for selection by using the differences in allele frequencies through time (or across space) to estimate the posterior odds (PO) of loci being under selection. For each locus, it also calculates the minimum false discovery rate (FDR) for selection as well (e.g., the probability that a given posterior probability of selection is likely to be a false positive). We considered loci with an FDR ≤0.05 and a log_10_PO ≥1 to be candidates for selection.

### Demographic Modeling with momi2 and Estimates of Temporal *N*_*e*_

Using momi2 v.2.1.19^35^, we tested among a series of demographic models to identify temporal fluctuations in *N*_*e*_ for each species. Specifically, we tested among fifteen demographic models: (1) no temporal change in *N*_*e*_, (2) a recent change in *N*_*e*_ between the historical and contemporary collection time points, (3) a historical change in *N*_*e*_ within 400 years prior to the historical collection time point, (4) an “ancient” change in *N*_*e*_ 10,000 - 1,000,000 years ago, (5) a recent and “ancient” change in *N*_*e*_, (6) a historical and “ancient” change in *N*_*e*_, (7) a recent and historical change in *N*_*e*_, (8) all three changes in *N*_*e*_, and (9-15) each of the previous seven scenarios as gradual (exponential) rather than abrupt (instantaneous) changes in *N*_*e*_ (SI Appendix, Fig. S9) We fit the models to (1) the contemporary SFS, (2) the historical SFS, and (3) both the historical and contemporary SFS. momi2 was chosen because it reliably reconstructs historical changes in *N*_*e*_, particularly for capture-based data^36^. For more information on model specification, see the SI Appendix, Supporting Methods. We fit all models to the observed SFS using the Truncated Newton (TNC) optimizer. The model with the lowest Akaike’s Information Criterion (AIC) was considered to be the most parsimonious and best model^80^. To estimate 95% CIs for the parameters of the best model, we generated 100 bootstrapped SFS from the observed SFS with momi2. The best demographic model was then fit to each bootstrapped SFS, and the 2.5th and 97.5th quantiles of the bootstrapped parameter estimates were calculated to create the 95% CI.

In addition to the demographic modeling, we used two methods to calculate temporal estimates of *N*_*e*_: (1) the allele frequency change method, by calculating the Jorde-Ryman modified temporal estimator^39^, and (2) the heterozygosity loss method, which applies the well-defined equation for estimating heterozygosity decline due to drift^41^ to calculate *N*_*e*_. Both estimates were generated using a dataset that was not LD-pruned. We used NeEstimator v.2.0.1^40^ to calculate the Jorde-Ryman estimates of *N*_*e*_, excluding loci with a minor allele frequency <0.05 and jackknife resampling across loci to calculate 95% CIs. For the heterozygosity loss *N*_*e*_ estimates, 95% CIs were generated from the bootstrapped *H*_*E*_ estimates.

## Data Availability Statement

Sequence and metadata are deposited in the NCBI SRA (project numbers: PRJNA998814, PRJNA998057, PRJNA998845, & PRJNA999299) and GEOME-DB (links: https://n2t.net/ark:/21547/FMB2, https://n2t.net/ark:/21547/FMH2, https://n2t.net/ark:/21547/FMR2, & https://n2t.net/ark:/21547/FMX2), respectively. Scripts for the results presented here are available at https://github.com/philippinespire/genetic_div_hamilo and archived on Zenodo (DOI:10.5281/zenodo.15085162).

## Acknowledgments

The authors thank the Municipality of Nasugbu (Batangas) and Bureau of Fisheries and Aquatic Resources for issuance of prior informed consent and sampling permits in the Philippines (Department of Agriculture Gratuitous Permit No. 0166-18), Angel Alcala for senior Philippine institutional administration through Silliman University-Angelo King Center for Research and Environmental Management, Jeffrey Williams for help in sampling and access to Smithsonian US Natural History Museum specimens (also facilitated by Diane Pitassy and other USNM curatorial staff), and Maddy Kenton and Ivan Lopez for help with extractions, library preparation, sequencing, and bioinformatics. This work was funded by a Rutgers EOAS Fellowship, a Rutgers SEBS Excellence Fellowship, and National Science Foundation nos OISE-1743711 and DEB-2343787. This research is based on work presented in the dissertation of René D. Clark^81^, but has not been published in its current format.

## Notes

https://doi.org/10.5281/zenodo.15085162

### Competing Interest Statement

The authors have declared no competing interest.

## References

1. S. Hoban et al., Global genetic diversity status and trends: Towards a suite of Essential Biodiversity Variables (EBVs) for genetic composition. Biol. Rev. 97, 1511–1538 (2022).

2. A. I. Thomson et al., Charting a course for genetic diversity in the UN Decade of Ocean Science. Evol. Appl. 14, 1497–1518 (2021).

3. D.M., Leigh, A. P. Hendry, E. Vázquez-Doínguez, V. L. Friesen, Estimated six per cent loss of genetic variation in wild populations since the Industrial Revolution. Evol. Appl. 12, 1505–1512 (2019).

4. M. Exposito-Alonso et al, Genetic diversity loss in the Anthropocene. Science 377, 1431–1435 (2022).

5. R. E. Shaw et al, Global meta-analysis shows action is needed to halt genetic diversity loss. Nature 636, 704–710 (2025).

6. A. Miraldo, An Anthropocene map of genetic diversity. Science 353, 15322–1535 (2016).

7. R. D. Clark et al., The practice and promise of temporal genomics for measuring evolutionary responses to global change. Mol. Ecol. Resour. 25, e13789 (2025).

8. K. E. Carpenter et al., One-third of reef-building corals face elevated extinction risk from climate change and local impacts. Science 321, 560–563 (2008).

9. M.I. O’Connor et al., Temperature control of larval dispersal and the implications for marine ecology, evolution, and conservation. Proc. Natl. Acad. Sci. U.S.A. 104, 1266–1271 (2007).

10. P. R. Teske, J. Sandoval-Castillo, E. van Sebille, J. Waters, L. B. Beheregaray, Oceanography promotes self-recruitment in a planktonic larval disperser. Sci. Rep. 6, 34205 (2016).

11. F. Marandel et al., Estimating effective population size of large marine populations, is it feasible? Fish Fish. 20, 189–198 (2019).

12. J. H. Steele, K. H. Brink, B. E. Scott, Comparison of marine and terrestrial ecosystems: Suggestions of an evolutionary perspective influenced by environmental variation. ICES J. Mar. Sci. 76, 50–59 (2019).

13. K. Y. Han et al., Genomic evidence for fisheries-induced evolution in Eastern Baltic cod. Sci. Adv. 11, eadr9889 (2025).

14. N. O. Therkildsen, E. E. Nielsen, D. P. Swain, J. S. Pedersen, Large effective population size and temporal genetic stability in Atlantic cod (Gadus morhua) in the southern Gulf of St. Lawrence. Can. J. Fish. Squat. Sci. 67, 1585–1595 (2010).

15. M. L. Pinsky, S. R. Palumbi, Meta-analysis reveals lower genetic diversity in overfished populations. Mol. Ecol. 2, 29–39 (2014).

16. M. L. Pinsky et al., Genomic stability through time despite decades of exploitation in cod on both sides of the Atlantic. Proc. Natl. Acad. Sci. U.S.A. 118, e2025453118 (2021).

17. A. D. Rogers et al. in The Blue Compendium: From Knowledge to Action for a Sustainable Ocean Economy 333–392 (Cham: Springer International Publishing, 2023).

18. K. R. Johnson, I. F. P. Owens, The Global Collection Group, A global approach for natural history museum collections. Science 379, 1192–1194 (2023).

19. D. G. Smith, J. T. Williams, The great Albatross Philippine expedition and its fishes. Mar. Fish. Rev. 61, 31–41 (1999).

20. R. V. Pagunsan, Mapping and narrating Philippine waters: Empire and science in the Albatross expedition to the US colony. Int. Rev. Environ. Hist. 4, 93–109 (2018).

21. Sparks, J. S. & Chakrabarty, P. A new species of ponyfish (Teleostei: Leognathidae: Photoplagios) from the Philippines. Copeia 3, 622–629 (2007).

22. M. E. Bloch, Naturgeschichte der Ausländischen Fische (Verfasser, 1795).

23. D. J. Woodland, S. Premcharoen, A. S. Cabanban in Species Identification Guide for Fishery Purposes. The Living Marine Resources of the Western Central Pacific. Bony Fishes Part 3 2792–2823 (FAO, Rome, 2001).

24. J. S. Sparks, P. V. Dunlap, W. L. Smith, Evolution and diversification of a sexually dimorphic luminescent system in ponyfishes (Teleostei: Leiognathidae), including diagnoses for two new genera. Cladistics 21, 305–327 (2005).

25. I. Asaad, C. J. Lundquist, M. V. Erdmann, M. J. Costello, Digital map of the Coral Triangle: An online atlas for marine biodiversity conservation. Earth Syst. Sci. Data Discuss., 1–19 (2018).

26. G. S. Jacinto, I. B. Velasquez, M. L. San Diego-McGlone, C. L. Villanoy, F. B. Siringan in The Environment in Asia Pacific Harbours 293–307 (Dordrecht: Springer Netherlands, 2006).

27. Vallejo B. M., Aloy, A. B., Ocampo, M., Conejar-Espedido, J., Manubag, L. M. in Impacts of Invasive Species on Coastal Environments: Coasts in Crisis 145–169 (Springer, 2019).

28. R. D. Lasco, E. Q. Javier, Lagunda de Bay: A case study for sustainable fisheries development. Trans. NAST PHL 39, 275–278 (2017).

29. A. Gnirke et al., Solution hybrid selection with ultra-long oligonucleotides for massively parallel targeted sequencing. Nat. Biotechnol. 27, 182–189 (2009).

30. E. L. Petrou et al., Intraspecific DNA contamination distorts subtle population structure in a marine fish: Decontamination of herring samples before restriction site-associated sequencing and its effects on population genetic statistics. Mol. Ecol. Resour. 19, 1131–1143 (2019).

31. D. H. Alexander, J. Novembre, K. Lange, Fast model-based estimation of ancestry in unrelated individuals. Genome Res. 19, 1655–1664 (2009).

32. G. Luikart, J.-M. Corneut, Estimating the effective number of breeders from heterozygote excess in progeny. Genet. 151, 1211–1216 (1999).

33. R. Frankham, Relationship of genetic variation to population size in wildlife. Conserv. Biol. 10, 1500–1508 (1996).

34. M. Foll, O. Gaggiotti, A genome-scan method to identify selected loci appropriate for both dominant and codominant markers: A Bayesian perspective. Genet. 180, 977–993 (2008).

35. J. Kamm, J. Terhost, R. Durbin, Y. S. Song, Efficiently inferring the demographic history of many populations with allele count data. J. Amer. Statist. Assoc. 115, 1472–1487 (2020).

36. B. N. Reid, M. L. Pinsky, Simulation-based evaluation of methods, data types, and temporal sampling schemes for detecting recent population declines. Integr. Comp. Biol. 62, 1849–1863 (2022).

37. W. B. Ludt, L. A. Rocha, Shifting seas: the impacts of Pleistocene sea-level fluctuations on the evolution of tropical marine taxa. J. Biogeogr. 42, 25–38 (2015).

38. A. C. Beichman, E. Huerta-Sanchez, K. E. Lohmueller, Using genomic data to infer historic population dynamics of nonmodel organisms. Annu. Rev. Ecol. Evol. Syst. 49, 433–456 (2018).

39. P. E. Jorde, N. Ryman, Unbiased estimator for genetic drift and effective population size. Genet. 177, 927–935 (2007).

40. C. Do et al., NeEstimator v2: Re-implementation of software for the estimation of contemporary effective population size (Ne) from genetic data. Mol. Ecol. Resour. 14, 209–214 (2014).

41. J. F. Crow, M. Kimura, An Introduction to Population Genetics Theory (Harper & Row, 1970).

42. R. S. Waples, The idiot’s guide to effective population size. Mol. Ecol., e17670 (2025).

43. L. Xu et al., Ecological connectivity of the Qiongzhou Strait: a case form orangefin ponyfish (Photopectoralis bindus) haplotype diversity and genetic structure. Front. Mar. Sci. 11, 1450142 (2024).

44. G. P. Jones, M. J. Milicich, M. J. Emslie, C. Lunow, Self-recruitment in a coral reef fish population. Nature 402, 802–804 (1999).

45. K. G. Sarakinis, et al. Strong philopatry in an estuarine-dependent fish. Ecol. Evol. 14, e10989 (2024).

46. L. H. Furness, et al. Historical collections of tropical marine mammals are an excellent resource for ancient DNA. Mol. Ecol. Resour., e70015 (2025).

47. G. Silvestre, R. Federizon, J. Munoz, D. Pauly, Over-exploitation of the demersal resources of Manila Bay and adjacent areas. Symposium on the Exploitation and Management of Marine Fishery Resources in Southeast Asia, 16–19 (1987).

48. S. López-Martínez, C. Morales-Caselles, J. Kadar, M. L. Rivas, Overview of global status of plastic presence in marine vertebrates. Glob. Change Biol. 27, 728–737 (2021).

49. D. M. Kummer, Measuring forest decline in the Philippines: An exercise in historiography. For. Conserv. Hist. 36, 185–189 (1992).

50. R. U. Briones, V. B. Ella, N. C. Bantayan, Hydrologic impact evaluation of land use and land cover change in Palico Watershed, Batangas, Philippines, using the SWAT model. Glob. J. Environ. Sci. Manag. 19, 96–107 (2016).

51. C. S. Rogers, Responses of coral reefs and reef organisms to sedimentation. Mar. Ecol. Prog. Ser. 62, 185–202 (1990).

52. D. M. Cabahug Jr., F. M. Ambi, S. O. Nisperos, T. C. Truzan Jr., Impact of community-based mangrove forestation to mangrove dependent families and to nearby coastal areas in Central Visayas: a case example. Mangroves of Asia and the Pacific: Status and Management (National Mangrove Committee and Natural Resources Management Center, 1986).

53. J. E. Limbo-Dizon, N. H. Dagamac, Assessment of coastal change detection on an urban coastline: A case study in metropolitan Manila. IOP Conf. Ser.: Earth Environ. Sci. 1165, 012015 (2023).

54. R. R. Diwa, C. C. Deocaris, L. P. Belo, River influx drives heavy metal pollution in Manila Bay, Philippines: An insight from multivariate analyses. Preprints doi: 10.20944/preprints202106.0470.v1 (2021).

55. H. O. Arceo, M. J. P. Velos, M. A. C. Nuñez, P.M. Aliño, Eds., The West Philippine Sea: State of the Coasts (University of the Philippines Marine Science Institute, 2024).

56. M. L. San Diego-McGlone, et al., Fish kills related to harmful algal bloom events in Southeast Asia. Sustainability 16, 10521 (2024).

57. F. X. D. Verdadero et al., Status of coral communities and reef-associated fish and invertebrates in Batangas and Northern Palawan. Manila J. Sci. 10, 101–114 (2017).

58. E. Karachaliou, C. Schmidt, E. de Greef, M. F. Docker, C. J. Garroway, Urbanisation is associated with reduced genetic diversity in marine fish populations. Mol. Ecol., e17711 (2025).

59. K. E. Carpenter, V. G. Springer, The center of the center of marine shore fish biodiversity: The Philippine Islands. Environ. Biol. Fishes 72, 467–480 (2005).

60. S. Díaz et al., Set ambitious goals for biodiversity and sustainability. Science 370, 411–413 (2020).

61. R. L. Roberts, “Whole genome sequencing century-old ethanol-preserved Philippine fishes,” thesis, Texas A&M University - Corpus Christi (2023); https://hdl.handle.net/1969.6/97716.

62. C. E. Bird et al., Philippines PIRE FQ GZ analysis pipeline, version 1.0.0, (2024); 10.5281/zenodo.13988543.

63. M. Vasimuddin, S. Misra, H. Li, S. Aluru in IEEE International Parallel and Distributed Processing Symposium (IPDPS) 314–324 (2019).

64. S. Dolenz et al., Unravelling reference bias in ancient DNA datasets. Bioinformatics 40, btae436 (2024).

65. A. Oliva, R. Tobler, A. Cooper, B. Llamas, Y. Souilmi, Systematic benchmark of ancient DNA read mapping. Brief. Bioinform. 22, bbab076 (2021).

66. W. Xu et al., An efficient pipeline for ancient DNA mapping and recovery of endogenous ancient DNA from whole-genome sequencing data. Ecol. Evol. 11, 390–401 (2020).

67. A. Bankevich et al., SPAdes: A new genome assembly algorithm and its applications to single-cell sequencing. J. Comput. Biol. 19, 455–477 (2012).

68. E. Garcia et al., Philippines PIRE SSL analysis pipeline, version 1.0.0, (2024); 10.5281/zenodo.13984548.

69. P. Danecek et al., Twelve years of SAMtools and BCFtools. GigaScience 10, giab008 (2021).

70. H. Jónsson, A. Ginolhac, M. Schubert, P. L. F. Johnson, L. Orlando, mapDamage2.0: Fast approximate Bayesian estimates of ancient DNA damage parameters. Bioinformatics 29, 1682–1684 (2013).

71. E. Garrison, G. Marth, Haplotype-based variant detection from short-read sequencing. 1207.3907 [genomics] (2012).

72. P. Danecek et al., The variant call format and VCFtools. Bioinformatics 27, 2156–2158 (2011).

73. S. Purcell et al., PLINK: A tool set for whole-genome association and population-based linkage analyses. Am. J. Hum. Genet. 81, 559–575 (2007).

74. R. M. Francis, pophelper:An R package and web app to analyse and visualize population structure. Mol. Ecol. Resour. 17, 27–32 (2017).

75. R Core Team, R: A Language and Environment for Statistical Computing (R Foundation for Statistical Computing, 2022). (2022).

76. J. Goudet, HIERFSTAT, a package for R to compute and test hierarchical F-statistics. Mol. Ecol. Notes 5, 184–186 (2005).

77. K. L. Korunes, K. Samuk, pixy: Unbiased estimation of nucleotide diversity and divergence in the presence of missing data. Mol. Ecol. Resour. 21, 1359–1368 (2021).

78. A. Canty, B. Ripley, boot: Bootstrap R (S-Plus) Functions, version 1.3.28 (2021); 10.32614/CRAN.package.boot.

79. R. N. Gutenkunst, R. D. Hernandez, S. H. Williamson, C. D. Bustamante, Inferring the joint demographic history of multiple populations from multidimensional SNP frequency data. PLoS Genet. 5, e1000695 (2009).

80. K. P. Burnham, D. R. Anderson, Model Selection and Multimodal Inference: A Practical Information-Theoretic Approach (Springer New York, 2002).

81. R. D. Clark, “Spatial and temporal patterns of adaptation and adaptive potential in a changing ocean,” thesis, Rutgers University (2023); 10.7282/t3-ycs5-z730.

