## Supplemental Methods and Figures for "Anthropocene genetic diversity loss in the marine tropics"

### Supporting Information Text

### Supporting Methods

**Capture bait design.** Twenty-four contemporary samples for each species were used to develop species-specific capture baits. Capture design samples were prepared using the single digest newRAD protocol, *sans* capture[1], and with modifications described in Roberts[2]. Briefly, total genomic DNA was digested with the *SbfI-HF* restriction enzyme (New England Biolabs), stubby biotinylated adapters with an internal barcode were ligated, and the libraries were sheared to approximately 300-500 bp using a Biorupter (Diagenode) sonicator. Streptavidin Dynabeads (ThermoFisher Scientific M-280) were used to enrich the libraries for DNA fragments with the biotinylated adapters. A second *Sbfl* digestion was performed to remove the biotin-Dynabead complex, and combinatorial dual indexed Illumina adapters were ligated using the KAPA Biosystems HyperPrep DNA kit. Libraries were normalized using the Biotium AccuBlue fluorescent dsDNA quantitation kit and pooled by species. Real-time qPCR on an Applied Biosystems StepOnePlus thermal cycler (ThermoFisher Scientific) was used to adjust the molarity of each library prior to combining into a super library. Paired end 2 x 150 bp sequencing was performed on an Illumina HiSeq 4000 at the Novogene Corporation Inc. (Sacramento, CA) to a target depth of 3 million reads per individual.

Reads were demultiplexed by Illumina index using the process_radtags function in STACKS[3]. We then deployed dDocentHPC[4], a modified fork of the dDocent bioinformatics pipeline[5] to trim, assemble, and map all reads. Briefly, adapter sequences, bases with a quality score <20, and reads less than 140 bp long (after trimming) were removed with Trimmomatic v.0.38[6], and *de novo* reference assemblies were created for each species using Rainbow v.2.0.4[7]. Reads were mapped to these *de novo* assemblies with BWA-mem2 v.2.2.1[8]. Improper read pairs and PCR clones were filtered with SAMtools v.1.9[9]. Variants were called with FreeBayes v.1.3.1[10].

To create a curated list of contigs that would be appropriate for bait creation, we applied the following filters to each *de novo* reference assembly with VCFtools v.0.1.14[11]: (1) filtered out contigs with abnormally low or high mean depth of coverage (<0.01 or >0.99 percentiles), as well as those with a particularly high coefficient of variation in mean depth of coverage across sites (>0.95 percentile), as they are unlikely to be reliably sequenced in all individuals downstream; (2) removed contigs with unusually high levels of heterozygosity (>90% of all sites); (3) removed contigs identified as paralogs with rad_haplotyper[12]; (4) filtered out contigs with repeat motifs as they may promote allelic dropout; (5) removed contigs with large stretches of missing data (>40 bp long), as they may have been improperly assembled; (6) removed contigs that are significantly longer (>95th quantile) or shorter (<5th quantile) than most, as they may either be improperly assembled or incomplete; (7) filtered out contigs with 2 or more indels of at least 3 nucleotides in length. After applying filters, 19,987 contigs were suitable for bait creation for *G. minuta*, while 32,765 contigs were suitable for *E. laterofenestra*.

Filtered *de novo* reference assemblies were then sent to Arbor Biosciences for capture bait design. Custom MYcroarray MYbaits kits were created for both species and designed so that each RAD locus (120 bp) was targeted by an average of three baits (i.e., 80 nt baits at an average of 3X tiling density). At most, one RAD locus was created per contig. A total of 19,998 baits were created for *G. minuta*, targeting 6,666 loci. For *E. laterofenestra,* a total of 13,658 baits were generated to target 4,688 loci. Presence or absence of variants was not a criteria in locus selection so as not to bias downstream estimates of genetic diversity.

**Extraction protocol, library preparation, and sequencing.** DNA extraction took place using the Qiagen Blood & Tissue Kit with the following modifications: (1) the volume of Proteinase K and Buffer ATL was doubled, (2) the volume of RNase A was doubled, (3) all centrifuge steps were performed 2x, and (4) each elution was stored separately. DNA was eluted (4x) with the Qiagen Buffer AE and visualized on 2% agarose gels to determine both quality and fragment length distribution.

Library preparation was performed by the Genomics Core Laboratory (GCL) at Texas A&M University - Corpus Christi. DNA extractions were subjected to a second round of gel electrophoresis and scored by smear pattern. For each individual, the elution with the highest ratio of high:low molecular weight fragments was used in subsequent reactions. Samples with molecular weight DNA <3000 bp were cleaned using Omega Biotek paramagnetic beads. We assessed DNA concentration with Biotium AccuClear kits (3 reactions/sample) on a SpectraMax M3 Reader and quality with a NanoDrop spectrophotometer. Capture libraries, for both historical and contemporary individuals, were prepared following Roberts[2] with the addition of a hybridization capture protocol[13]. For information on the capture bait design, see the "Capture Bait Design" section above. The KAPA Biosystems HyperPlus DNA prep kit was used to randomly fragment genomic DNA enzymatically prior to ligation with dual combinatorially indexed Illumina adapters. Ligated samples were normalized using the Biotium AccuClear dsDNA quantitation kit and combined into four pools by species and sampling time before capture following the MYcroarray MYbaits protocol and size-selection (400-650 bp) using a Blue Pippin (Sage Science). Library fragment lengths were quantified using an Advanced Analytical Fragment Analyzer. The pools were normalized using real-time qPCR with the KAPA Library Quantification Kit on an Applied Biosystems StepOnePlus ThermoCycler (ThermoFisher Scientific) and combined into a superpool.

To avoid contamination, neither extraction or library preparation of contemporary and historical samples were performed on the same day, stored on the same plates, or unsealed at the same time. All countertops, equipment, and tools were bleached before and after the protocols were executed. Additionally, DNA extraction, pre-PCR, and post-PCR sample handling occurred in separate, designated areas in the GCL.

**Data processing and filtering.** Raw reads were quality assessed with FastQC v.0.11.9[14] and trimmed with *fastp* v.0.20.0[15] to remove adapters, poly-G tails, unpaired reads, low complexity reads, bases with a quality score <20 from the 3' ends, and reads that were less than 33 bp long (after trimming). Reads were then de-duplicated and sorted with *clumpify* from the BBTools suite of bioinformatic tools[16] and trimmed again with *fastp* to remove bases with a quality score <20 at the 5' read ends. Next, contaminated reads that contained exogenous DNA were identified and filtered with FastQ Screen v.0.14.1[17], by using Bowtie2 v.2.4.1[18] to map all reads to a database of all known sequencing oligonucleotides and adapters; 24,979 bacterial; 1,046 protist; 42,368 viral; and 80 fungal genomes, in addition to a subset of potential contaminant agents from the laboratories where library preparation occurred (two coral genomes, one bird genome, one plant genome, and the human genome [GRCh38]). Reads that mapped to any of the contaminant genomes were removed. Finally, reads were re-paired using *repair.sh* from BBTools.

**SNP and individual filtering.** Filtering with VCFtools v.0.1.14[11] removed the following: (1) SNPs with a quality score <100, (2) SNPs missing in >45% of individuals, (4) SNPs with a mean depth <5 or >400 across all individuals, (5) SNPs with a quality/depth ratio <0.2, (6) SNPs where the difference in the ratio of mean mapping quality between reference and alternate alleles was >0.25, (7) SNPs with a mean allele balance <0.375 or >0.625 across all individuals, (8) SNPs where either the reference or alternate allele (or both) was supported only by reads that were improperly paired, (9) SNPs out of Hardy-Weinberg proportions (HWP) within a time point. SNPs were pruned for linkage disequilibrium (LD; $R^2$ >0.5) using PLINK v.1.9[19]. Additionally, individuals with >50% missing data were removed.

Prior to filtering for Hardy-Weinberg equilibrium, population structure was assessed with PCA and ADMIXTURE (see below for more details). During this step, if cryptic species were identified, the species delineation was incorporated into the population identification during HWP filtering, so that SNPs would not be considered out of HWP due to cryptic structure within a population. Populations were considered to be composed of two or more species if (1) ADMIXTURE assigned individuals entirely to separate genetic clusters and (2) the pairwise $F_{ST}$ value between those genetic clusters was >0.35. For *G. minuta*, we found that ADMIXTURE assigned ~50% of both the historical and contemporary populations to one genetic cluster and the remaining 50% to a separate cluster (pairwise $F_{ST}$ between clusters was 0.660 for historical individuals and 0.694 for contemporary individuals). We recovered and blasted the COI sequences for several individuals in both clusters. One cluster matched against *G. minuta* COI sequences with high sequence similarity and confidence, while the other cluster did not (and instead had higher percent matches to *Gazza achlamys* or other species in the *Gazza* genus). For analyses, we retained only those individuals in the putative *G. minuta* cluster (85 individuals in total: 19 historical, 66 contemporary). For *E. laterofenestra*, we found that ADMIXTURE assigned ~50% of the historical population to one genetic cluster and the remaining 50% (along with almost the entire contemporary population) to a separate cluster (pairwise $F_{ST}$ between clusters was 0.492 for the historical individuals). Subsequent morphological and genetic identification (via barcoding of several individuals from each cluster) confirmed that the second cluster was *Equulites laterofenestra*, while the first (comprised of mainly historical individuals) was likely a separate species in the *Equulites* genus (possibly *Equulites leuciscus*). Importantly, *E. laterofenestra* was not described until 2007[20], which may have contributed to the presence of mixed species in the *Equulites* lot. For analyses, we retained only those individuals in the putative *E. laterofenestra* cluster (105 individuals in total: 11 historical, 94 contemporary).

Finally, to control for contamination that may have occurred during sample storage or DNA library preparation, we calculated within-individual heterozygosity (the proportion of heterozygous loci within individuals) with VCFtools and removed any samples whose individual heterozygosity was three standard deviations above the species mean[21]. This removed 4 fish for *G. minuta* and 2 fish for *E. laterofenestra*. Notably, historical libraries were characterized by distinctly shorter sequences than contemporary libraries due to DNA degradation, and there were no indications of cross contamination between historical and contemporary samples.

**Historical DNA quality control.** Historical damage patterns were quantified with mapDamage2 v.2.2.1[22] and base quality scores were recalibrated to account for base-calling errors characteristic to historical DNA degradation (SI Appendix, Fig. S2). We checked for issues known to plague data from historical DNA, including: (1) reference bias[23,24], (2) heterozygosity-depth relationships that may bias temporal trends in genetic diversity due to variation in read depth across time points[25,26], and (3) unusual patterns in missing genotype calls. First, to test for reference bias, we examined allele balance and the read depth ratio. In general, we expect heterozygous genotypes to have an allele balance of 50:50, where 50% of the reads represent the reference allele and 50% of the reads represent the alternate allele. If reference bias is an issue, we would instead expect reads with the reference allele to map more frequently, and be more common, than those with alternate alleles, and thus would expect to see average allele balances <50%. This would be especially true in the historical populations, as these individuals are theoretically more diverged from the reference genome (that was made from contemporary individuals only) than those in the contemporary populations. Similarly, in datasets with reference bias issues, we would also expect to see a lower homozygous:heterozygous read depth ratio between genotypes at a given locus (as half the homozygous reads, those with alternate alleles, would not map as well).

We found no evidence of reference bias in either of our species: allele balance was centered around 50:50 (Fig. S3 a and b) and read depth ratio was approximately 1 (Fig. S3 c and d) in all historical and contemporary populations. Second, we calculated heterozygosity-depth relationships across all time points/populations, by quantifying read depth distributions in both historical and contemporary populations. If read depth differs dramatically across time points, samples with abnormally low read depth may underestimate heterozygosity or genetic diversity, as they have lower power for accurately calling heterozygote genotypes. In our data, a read depth of ~5-10X gave sufficient power to consistently and confidently call heterozygote genotypes (Fig. S3 e and f). The genome-wide average read depth was higher than this threshold in both species and at both time points (Tables S1 and S2). In addition, the genome-wide average read depth was higher than the more conservative 15-20X minimum read depth[25] suggested for avoiding these issues. Finally, we found no biased patterns in missing genotype calls in either of our species (Fig. S3 g and h). Tables S1 and S2 show a breakdown of total number of reads, mapped reads, mean read length, mean depth, and retained SNPs/individuals for both species and time points.

**momi2 model design.** We fit parameters for our demographic models to the observed SFS from the "all site" datasets and estimated the likelihood of our data under each model. The total number of sites was estimated as the number of loci in the "all sites" dataset from which the SFS was made. Parameters estimated from the models included: (1) ancient $N_e$ prior to the onset of the ancient change in $N_e$, (2) post-ancient $N_e$ prior to the onset of the historical change in $N_e$, (3) historical $N_e$ prior to onset of the recent change in $N_e$, (4) contemporary $N_e$ at the time of contemporary sampling, (5) the time of the ancient change in $N_e$, (6) the time of the historical change in $N_e$, and (7) the time of the recent change in $N_e$. For the models with gradual change in $N_e$, the rate of the size change was determined within the model and estimated from both the time of decline and the two $N_e$ parameters (e.g. for the recent population size change, the time of the size change and the modern and historical $N_e$). Importantly, when modeling population size change we did not explicitly force a decrease in population size, just that a change occurred (in either direction). For both species, the generation length was estimated to be 1.39 years, using the R package FishLife v.3.0.0[27]. For all models the mutation rate was set to $3.5 \times 10^{-9}$/site/generation, consistent with reported mutation rates for marine fishes.[28,29] As momi2[30] assumes a tree-like population history, we specified the temporal data (historical population) as a "leaf" from a branch originating at its corresponding contemporary sampling time (as applicable).

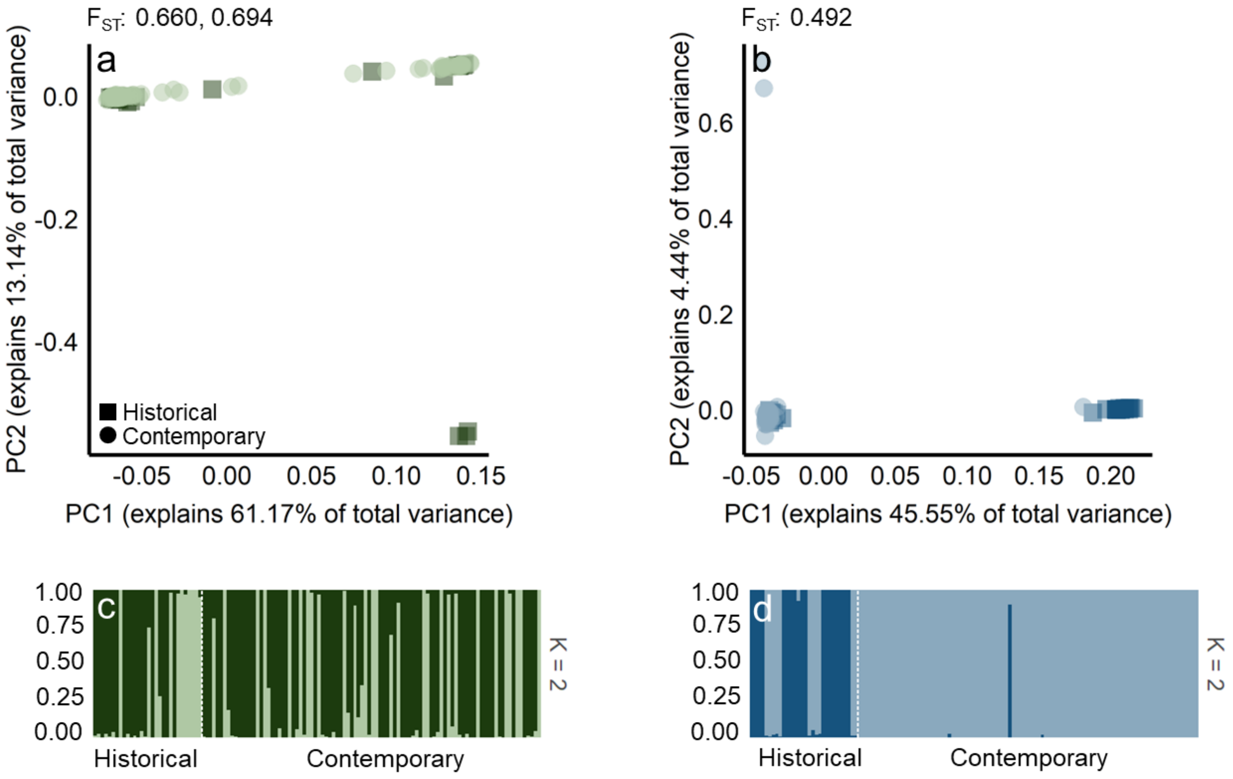


**Fig. S1.** PCAs (a: *G. minuta*, b: *E. laterofenestra*) & ADMIXTURE analyses manually forcing K = 2 (c: *G. minuta*, d: *E. laterofenestra*). For both PCAs, historical individuals are represented by dark squares and contemporary individuals are represented with lighter circles. For both ADMIXTURE plots, each putative population is represented by a unique color, individuals are grouped by time point (divided by the dashed vertical lines), and the proportion of assignment is represented by the vertical axis on the left. K=2 was best supported for both species. For all figures, cryptic species are included (e.g., individuals putatively identified as belonging to a cryptic species have not been removed prior to these analyses). High heterozygosity individuals were also not removed prior to analyses. Pairwise $F_{ST}$ across cryptic species but within time points is reported above each PCA (Historical, Contemporary). For *E. laterofenestra*, only historical pairwise $F_{ST}$ is reported.

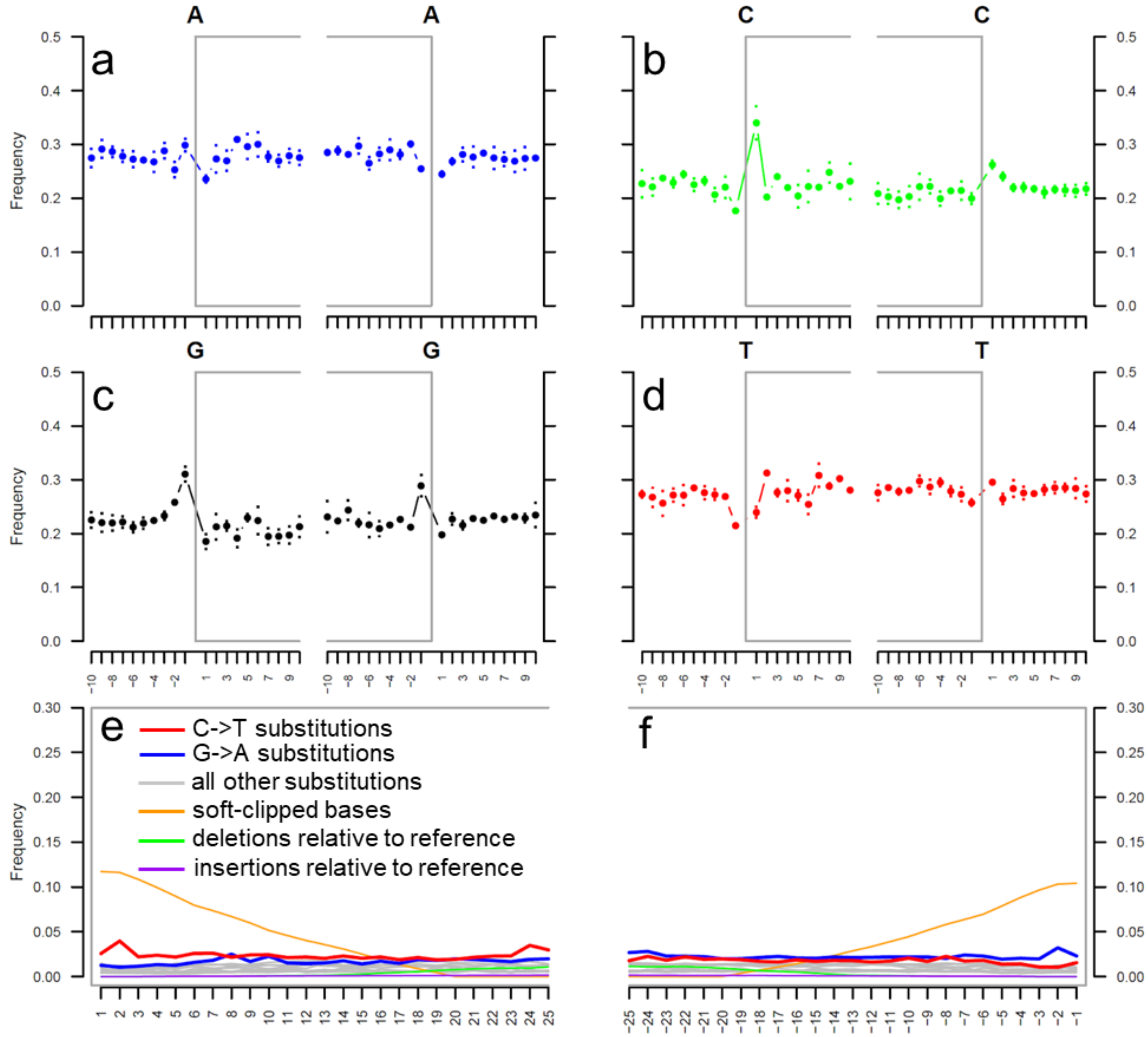


**Fig. S2.** Representative plots of historical DNA fragmentation and mis-incorporation patterns. From sequencing read data from *G. minuta* individual AHam-001. Fragmentation patterns; the base frequency outside and inside the read alignment (depicted by the position of the gray boxes for nucleotides (a) A, (b) C, (c) G, and (d) T. Mis-incorporation patterns; (e) elevated 5' end C->T substitution patterns (represented by the red line) and (f) elevated 3' end G->A substitution patterns (represented by the blue line). For the mis-incorporation pattern plots, the X-axes represent the nucleotide position relative to the 5' end (e) or 3' end (f) of each read and the orange lines represent the frequency of soft-clipped bases. Plots generated by MapDamage v.2.2.1[21]. All other historical individuals show similar patterns.

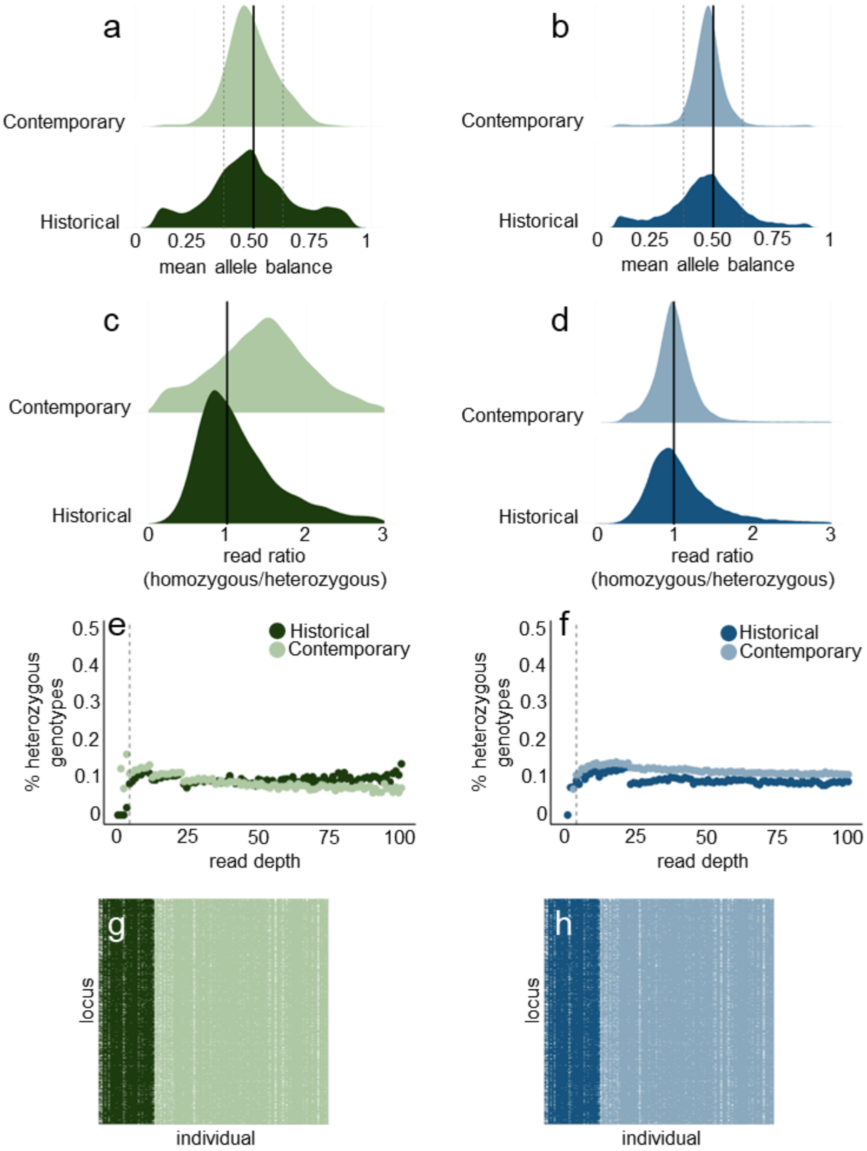


**Fig. S3.** Plots of genotype quality. Density plot of average allele balance across individuals per locus for (a) *G. minuta* and (b) *E. laterofenestra*. For both, vertical dashed lines indicate cut-offs for allele balance filtering. Density plot of read depth ratio for (c) *G. minuta* and (d) *E. laterofenestra*. Read depth ratio is calculated on a per locus basis and equals the average read count for all homozygous genotypes divided by the average read count for all heterozygous genotypes. Proportion of all genotypes that were heterozygous at any given read depth for (e) *G. minuta* and (f) *E. laterofenestra*. For both, vertical dashed lines indicate cut-offs for mean read depth filtering. Heat map of missing genotypes per individuals retained after filtering for (g) *G. minuta* and (h) *E. laterofenestra*. For both heat maps, rows represent loci, columns represent individuals. Cells are colored if a genotype was called and empty (white) if the genotype was missing. For all plots, darker colors represent historical individuals and lighter colors represent contemporary individuals. All genotyped individuals are included.

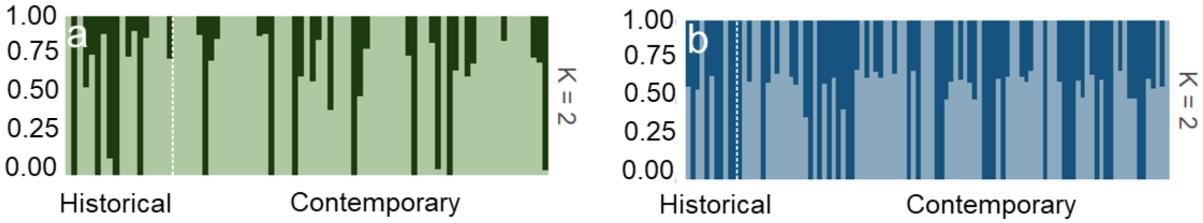


**Fig. S4.** ADMIXTURE plots manually forcing K = 2 after removing cryptic species and high heterozygosity individuals for (a) *G. minuta* and (b) *E. laterofenestra*. For both figures, each putative population is represented by a unique color, individuals are grouped by time point (divided by the dashed vertical lines), and the proportion of assignment is represented by the vertical axis on the left. K=1 was best supported for both species.

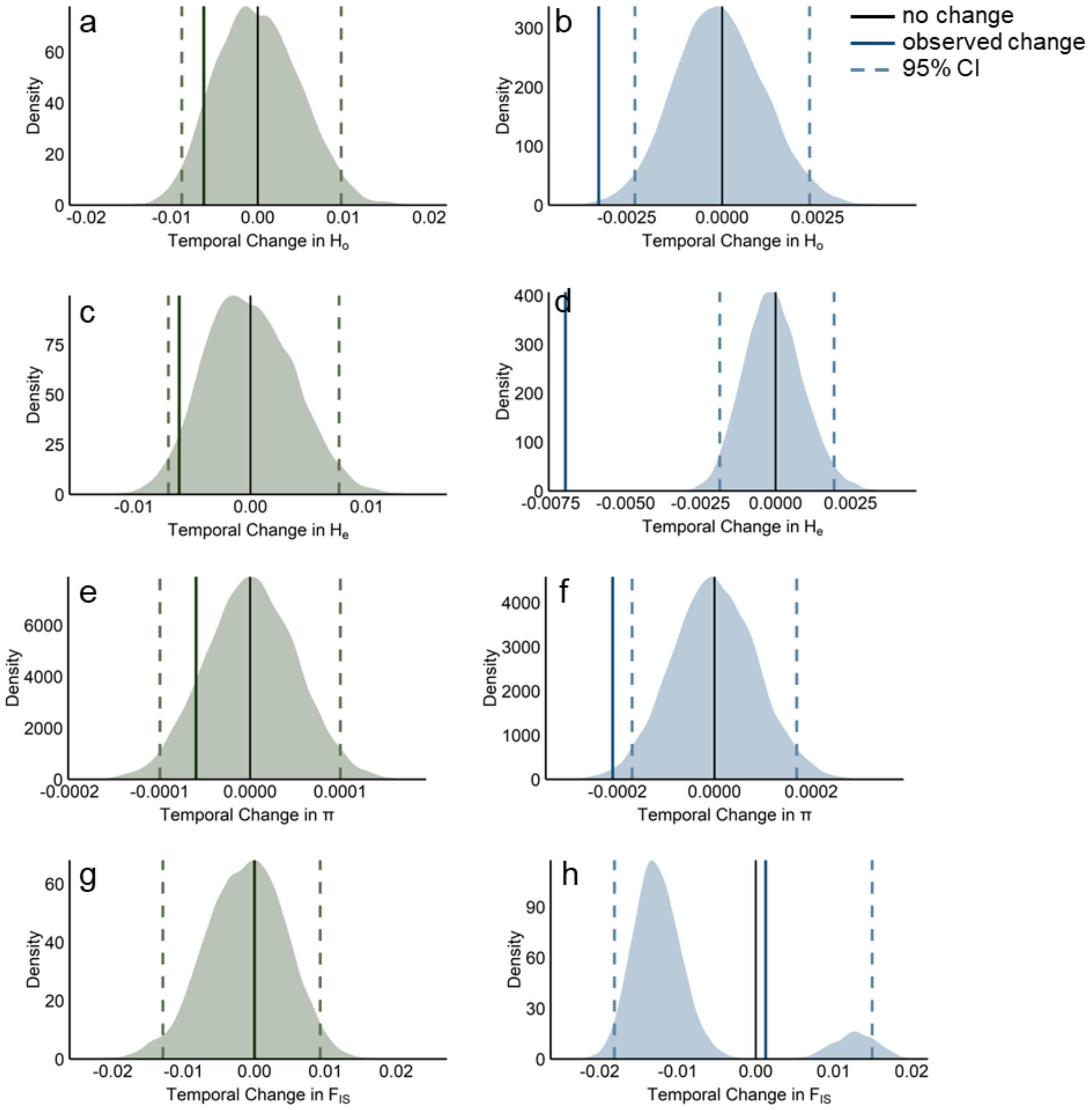


**Fig. S5.** Observed vs. expected changes in diversity under a null model of no change through time. The shaded area under the curve represents the null model changes through time in the test statistic for 10,000 permuted datasets. Black lines at zero mark no change through time. Solid colored lines represent the observed change in the real datasets, while dashed colored lines represent the 95% CI of null model change. (a) changes in $H_O$ for *G. minuta*; (b) changes in $H_O$ for *E. laterofenestra*; (c) changes in $H_E$ for *G. minuta*; (d) changes in $H_E$ for *E. laterofenestra*; (e) changes in π for *G. minuta*; (f) changes in π for *E. laterofenestra*; (g) changes in $F_{IS}$ for *G. minuta*, with the line for observed change directly over zero; (h) changes in $F_{IS}$ for *E. laterofenestra*.

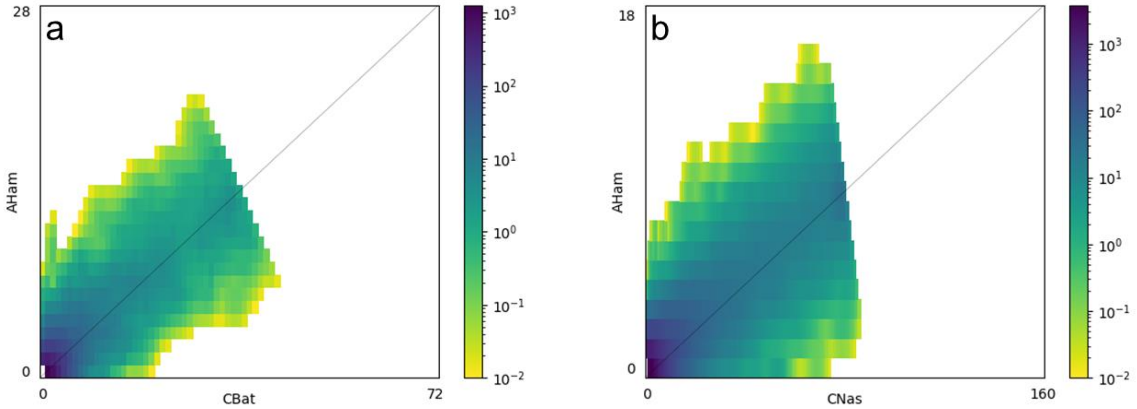


**Fig. S6.** 2-dimensional site frequency spectra (SFS) for (a) *G. minuta* and (b) *E. laterofenestra*. Allele frequencies (counts) for historical populations are on the y-axis, while those for contemporary populations are on the x-axis. Black diagonal line represents SNPs that have the same allele frequency in both time points. Darker colors represent higher numbers of SNPs.

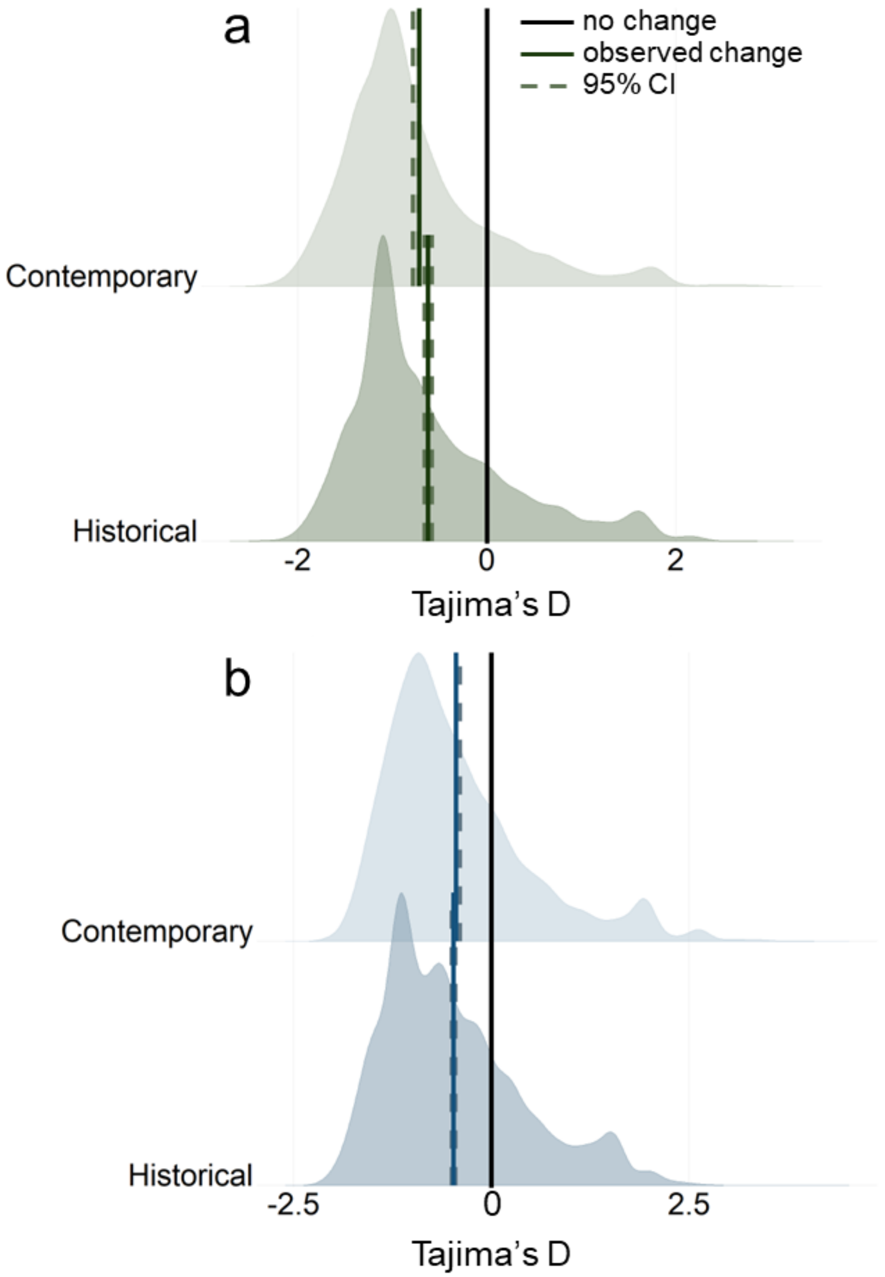


**Fig. S7.** Distribution of Tajima's *D* in historical and contemporary populations of (a) *G. minuta* and (b) *E. laterofenestra*. Black solid line at zero represents Tajima's *D* expected if populations are at mutation-drift equilibrium. Solid colored lines represent observed means and dashed colored lines represent the 95% CIs around the mean.

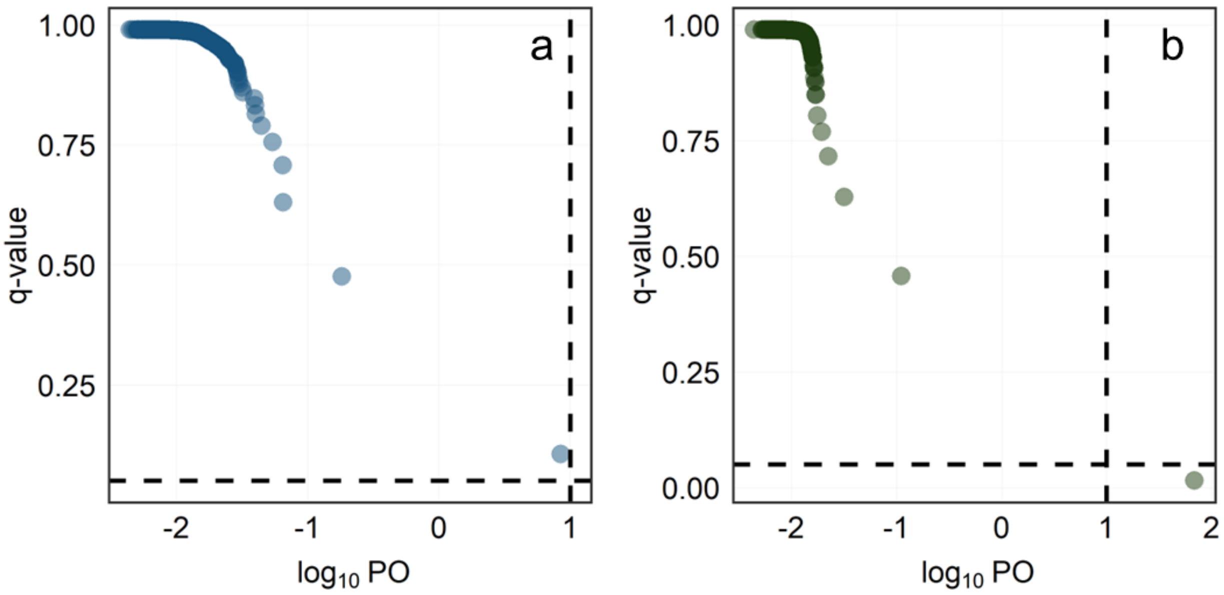


**Fig. S8.** BayeScan results for (a) *G. minuta* and (b) *E. laterofenestra*. For each SNP, the $\log_{10}$ posterior odds (PO) for the model including selection are on the x-axis and q-values (False Discovery Rates, FDR) are on the y-axis. The horizontal black dashed line represents the FDR significance threshold of 0.05. The vertical black dashed line at 1 represents the $\log_{10}$PO cut-off for "strong" evidence of selection. SNPs with an FDR $\leq 0.05$ and a $\log_{10}\text{PO} \geq 1$ are considered outliers and candidates for selection.

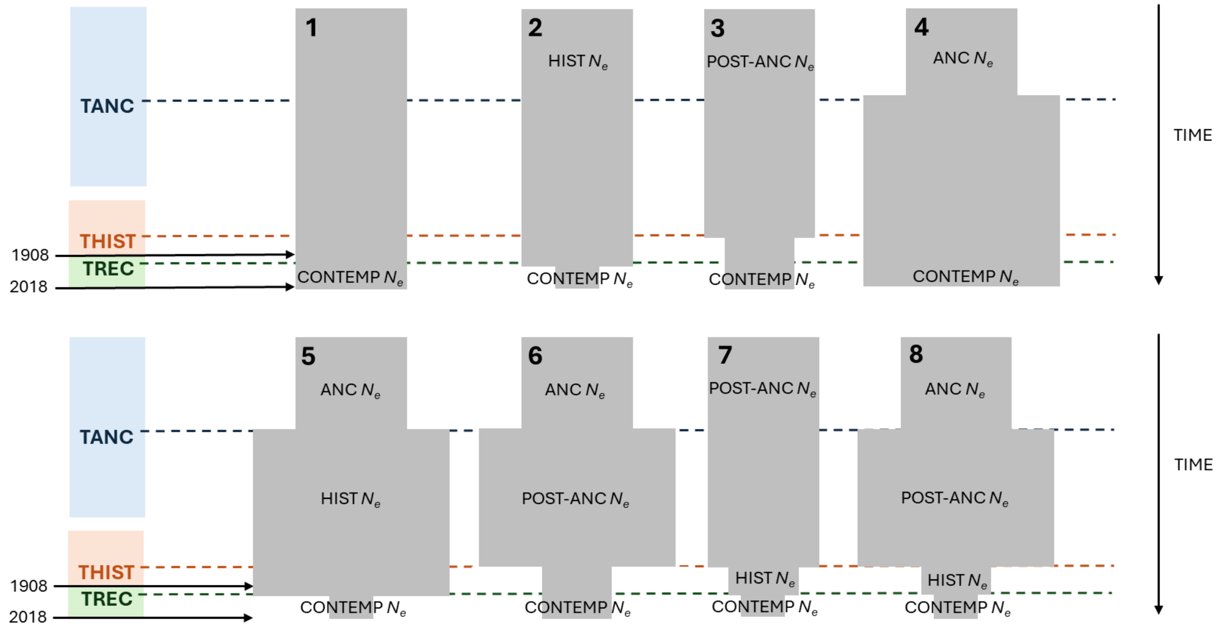


**Fig. S9.** Conceptual diagrams of demographic models. TANC represents the timing of the ancient change in $N_e$, THIST represents the timing of the historical change in $N_e$ , and TREC represents the timing of the recent change in $N_e$. Shaded boxes represent the possible timeframe during which these changes in $N_e$ could occur. ANC $N_e$ represents $N_e$ prior to the onset of the ancient population size change, POST-ANC $N_e$ represents $N_e$ prior to the onset of the historical population size change, HIST $N_e$ represents $N_e$ prior to the onset of the recent population size change, and CONTEMP $N_e$ represents $N_e$ of the contemporary population. Numbering corresponds to numbering of the different modeling scenarios in the methods section of the main text.(see methods section in the main text).

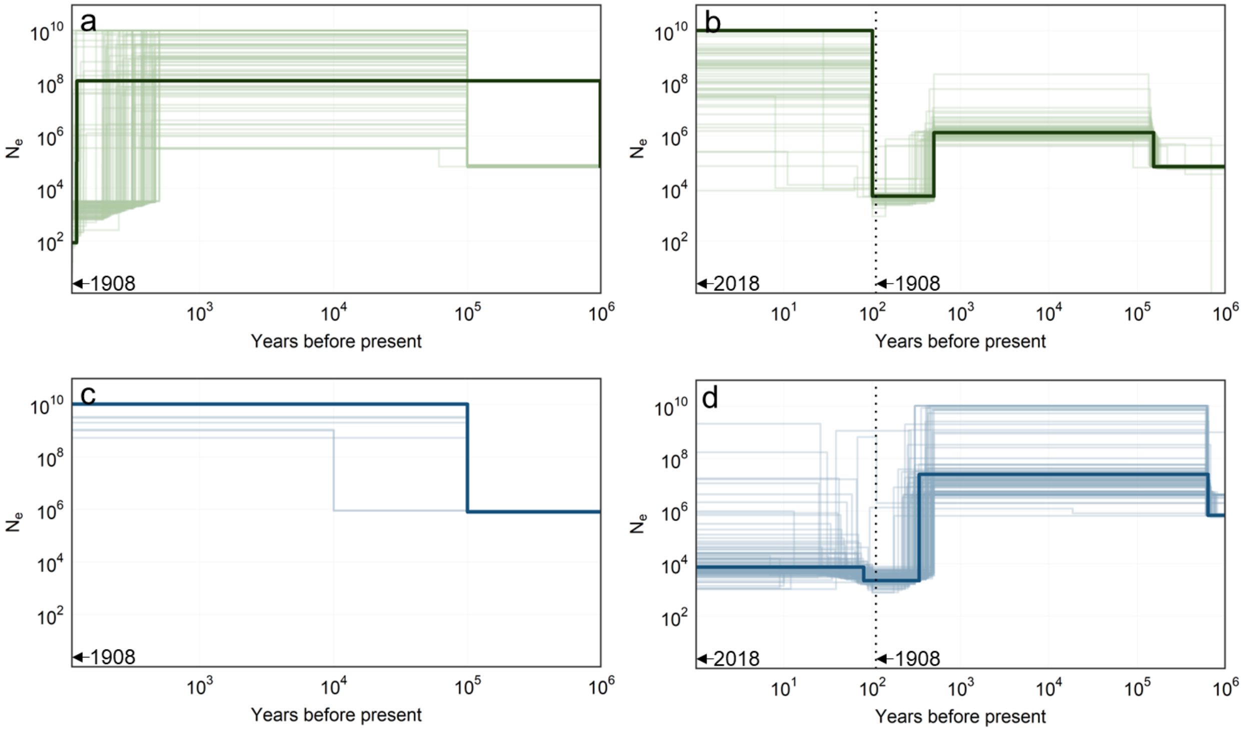


**Fig. S10.** Plots of $N_e$ through time with historical or temporal data. Both axes are depicted in log scale. For all figures, the solid dark line represents estimated $N_e$ from the most parsimonious model fit to the observed SFS, while lighter lines represent estimated $N_e$ from the most parsimonious model fit to 100 bootstrapped SFS (each line represents one bootstrap). (a) $N_e$ through time for *G. minuta* with models fit to the historical SFS only; (b) $N_e$ through time for *G. minuta* with models fit to the historical and contemporary SFS; (c) $N_e$ through time for *E. laterofenestra* with models fit to the historical SFS only; (d) $N_e$ through time for *E. laterofenestra* with models fit to the historical and contemporary SFS. For (b) and (d), the dashed vertical line indicates the 1908 time point. Estimates were generated using momi2; changes in $N_e$ were modeled as instantaneous growth (or decline).

**Table S1.** Sequencing statistics for all historical samples. Sample names are given in column one, species names in column two (*Gmi* = *G. minuta*, *Ela* = *E. laterofenestra*), the NMNH lot number in column three, and the year the sample was originally collected in column four. Column five lists the number of raw Illumina reads per sample. Column six gives the number of reads that mapped to the reference genome and passed filtering. Column seven lists the mean read length for mapped reads, while column eight gives the average read depth for each SNP used in most downstream analyses. Column nine gives the number of SNPs with missing genotypes for each individual. Column ten indicates whether or not the individual was used in the final analyses. An asterisk in the species column indicates the individual was identified as belonging to a cryptic species. HH in parentheses in the included column indicates the sample was not included in the final analyses as its individual heterozygosity was three standard deviations above the species mean.

| Individual | Species | NMNH Lot | Year | Raw Reads | Mapped Reads | Mean Read Length | Mean SNP Depth | Missing SNPs | Included |
|---|---|---|---|---|---|---|---|---|---|
| AHam-001 | *Gmi* | 191914 | 1908 | 19,962,372 | 392,786 | 130.2 | – | – | N |
| AHam-002 | *Gmi* | 191914 | 1908 | 27,486,820 | 888,832 | 137.0 | 9.3 | 3,537 | Y |
| AHam-003 | *Gmi* | 191914 | 1908 | 7,479,940 | 170,338 | 132.0 | 9.2 | 3,978 | Y |
| AHam-004 | *Gmi* | 191914 | 1908 | 22,009,610 | 680,670 | 131.8 | 51.3 | 15 | Y |
| AHam-005 | *Gmi* | 191914 | 1908 | 18,948,646 | 437,986 | 131.9 | – | – | N |
| AHam-006 | *Gmi* | 191914 | 1908 | 33,288,364 | 53,425 | 121.3 | – | – | N |
| AHam-007 | *Gmi* | 191914 | 1908 | 55,765,468 | 103,933 | 124.2 | – | – | N |
| AHam-008 | *Gmi* | 191914 | 1908 | 35,740,590 | 177,279 | 121.2 | – | – | N |
| AHam-009 | *Gmi* | 191914 | 1908 | 29,729,862 | 223,825 | 118.1 | – | – | N |
| AHam-010 | *Gmi* | 191914 | 1908 | 37,382,828 | 411,174 | 132.4 | 22.4 | 1,335 | Y |
| AHam-011 | *Gmi* | 191914 | 1908 | 20,801,260 | 644,597 | 127.8 | 39.8 | 36 | Y |
| AHam-012 | *Gmi* | 191914 | 1908 | 24,646,988 | 560,795 | 129.3 | – | – | N |
| AHam-013 | *Gmi* | 191914 | 1908 | 31,647,726 | 1,008,158 | 135.7 | 15.9 | 1,785 | Y |
| AHam-014 | *Gmi* | 191914 | 1908 | 19,663,402 | 592,388 | 122.7 | 43.7 | 31 | Y |
| AHam-015 | *Gmi* | 191914 | 1908 | 22,044,942 | 39,359 | 124.5 | – | – | N |
| AHam-016 | *Gmi** | 191914 | 1908 | 55,711,112 | 534,926 | 125.8 | 39.2 | 141 | N |
| AHam-017 | *Gmi* | 191914 | 1908 | 23,266,014 | 579,395 | 133.2 | 44.3 | 8 | Y |
| AHam-018 | *Gmi* | 191914 | 1908 | 11,250,450 | 269,345 | 124.4 | 22.3 | 296 | Y |
| AHam-019 | *Gmi* | 191914 | 1908 | 20,741,152 | 436,291 | 129.6 | – | – | N |
| AHam-020 | *Gmi* | 191914 | 1908 | 26,331,328 | 639,643 | 131.1 | – | – | N |
| AHam-021 | *Gmi* | 191914 | 1908 | 25,633,276 | 470,065 | 127.2 | – | – | N |

| | | | | | | | | | |
|---|---|---|---|---|---|---|---|---|---|
| AHam-022 | *Gmi* | 191914 | 1908 | 15,491,866 | 319,680 | 127.9 | – | – | N |
| AHam-023 | *Gmi* | 191914 | 1908 | 23,508,562 | 596,186 | 135.3 | – | – | N |
| AHam-024 | *Gmi* | 191914 | 1908 | 38,450,202 | 146,569 | 120.8 | – | – | N |
| AHam-025 | *Gmi* | 191914 | 1908 | 13,733,094 | 290,520 | 128.1 | – | – | N |
| AHam-026 | *Gmi* | 191914 | 1908 | 16,457,806 | 484,518 | 132.4 | 43.1 | 26 | Y |
| AHam-027 | *Gmi* | 191914 | 1908 | 14,662,256 | 425,809 | 133.3 | 33.6 | 71 | Y |
| AHam-028 | *Gmi* | 191914 | 1908 | 14,907,270 | 365,767 | 128.0 | 26.7 | 122 | Y |
| AHam-029 | *Gmi* | 191914 | 1908 | 11,809,976 | 235,003 | 125.1 | 14.8 | 2,292 | Y |
| AHam-030 | *Gmi* | 191914 | 1908 | 10,262,544 | 258,224 | 126.2 | 19.3 | 470 | Y |
| AHam-031 | *Gmi** | 191914 | 1908 | 53,310,460 | 719,325 | 120.8 | 51.4 | 27 | N |
| AHam-032 | *Gmi* | 191914 | 1908 | 17,515,428 | 456,970 | 125.0 | 31.0 | 71 | Y |
| AHam-033 | *Gmi** | 191914 | 1908 | 35,459,446 | 393,309 | 130.0 | 34.4 | 234 | N |
| AHam-034 | *Gmi* | 191914 | 1908 | 36,326,884 | 86,729 | 123.2 | – | – | N |
| AHam-035 | *Gmi* | 191914 | 1908 | 28,955,304 | 722,033 | 124.2 | 52.9 | 8 | N (HH) |
| AHam-036 | *Gmi* | 191914 | 1908 | 26,736,118 | 157,827 | 116.1 | – | – | N |
| AHam-037 | *Gmi* | 191914 | 1908 | 53,380,012 | 1,575,662 | 131.8 | 90.4 | 0 | Y |
| AHam-038 | *Gmi* | 191914 | 1908 | 11,204,382 | 201,240 | 121.6 | – | – | N |
| AHam-039 | *Gmi* | 191914 | 1908 | 29,311,940 | 1,014,883 | 135.1 | 70.4 | 4 | Y |
| AHam-040 | *Gmi* | 191914 | 1908 | 44,011,124 | 133,614 | 117.2 | – | – | N |
| AHam-041 | *Gmi** | 191914 | 1908 | 3,205,488 | 186,891 | 125.3 | 15.7 | 1,431 | N |
| AHam-042 | *Gmi* | 191914 | 1908 | 14,267,520 | 360,438 | 121.9 | 28.2 | 101 | Y |
| AHam-043 | *Gmi* | 191914 | 1908 | 25,119,394 | 145,081 | 118.9 | – | – | N |
| AHam-044 | *Gmi* | 191914 | 1908 | 36,998,132 | 179,935 | 119.1 | – | – | N |
| AHam-045 | *Gmi* | 191914 | 1908 | 53,954,062 | 324,928 | 118.4 | – | – | N |
| AHam-046 | *Gmi* | 191914 | 1908 | 39,625,902 | 229,546 | 119.6 | – | – | N |
| AHam-047 | *Gmi* | 191914 | 1908 | 38,874,850 | 72,868 | 124.2 | – | – | N |
| AHam-048 | *Gmi* | 191914 | 1908 | 51,211,728 | 40,614 | 120.2 | – | – | N |
| AHam-049 | *Gmi* | 191914 | 1908 | 26,310,002 | 38,396 | 124.5 | – | – | N |
| AHam-050 | *Gmi* | 191914 | 1908 | 15,274,752 | 286,610 | 126.1 | – | – | N |
| AHam-051 | *Gmi* | 191914 | 1908 | 27,866,720 | 157,263 | 126.4 | – | – | N |
| AHam-052 | *Gmi* | 191914 | 1908 | 17,013,052 | 418,582 | 135.7 | – | – | N |
| AHam-053 | *Gmi* | 191914 | 1908 | 17,968,756 | 130,773 | 114.0 | – | – | N |

| | | | | | | | | | |
|---|---|---|---|---|---|---|---|---|---|
| AHam-054 | *Gmi* | 191914 | 1908 | 14,657,114 | 303,434 | 126.6 | – | – | N |
| AHam-055 | *Gmi** | 191914 | 1908 | 35,813,496 | 210,756 | 126.7 | 19.1 | 898 | N |
| AHam-056 | *Gmi* | 191914 | 1908 | 26,663,096 | 194,349 | 120.3 | – | – | N |
| AHam-057 | *Gmi* | 191914 | 1908 | 46,030,520 | 205,248 | 121.9 | – | – | N |
| AHam-058 | *Gmi* | 191914 | 1908 | 29,824,562 | 184,127 | 116.2 | – | – | N |
| AHam-059 | *Gmi* | 191914 | 1908 | 38,001,662 | 30,913 | 122.8 | – | – | N |
| AHam-060 | *Gmi** | 191914 | 1908 | 57,804,462 | 420,699 | 124.5 | 34.4 | 211 | N |
| AHam-061 | *Gmi* | 191914 | 1908 | 33,551,448 | 77,274 | 119.4 | – | – | N |
| AHam-062 | *Gmi* | 191914 | 1908 | 14,654,172 | 379,497 | 129.9 | – | – | N |
| AHam-063 | *Gmi* | 191914 | 1908 | 27,033,802 | 175,543 | 114.0 | – | – | N |
| AHam-064 | *Gmi* | 191914 | 1908 | 28,198,468 | 190,048 | 121.0 | – | – | N |
| AHam-065 | *Gmi** | 191914 | 1908 | 128,000,000 | 514,383 | 124.0 | 36.8 | 365 | N |
| AHam-066 | *Gmi* | 191914 | 1908 | 23,741,762 | 142,617 | 119.4 | – | – | N |
| AHam-067 | *Gmi* | 191914 | 1908 | 66,208,928 | 118,979 | 123.7 | – | – | N |
| AHam-068 | *Gmi* | 191914 | 1908 | 39,926,112 | 108,811 | 121.8 | – | – | N |
| AHam-069 | *Gmi* | 191914 | 1908 | 37,675,422 | 56,089 | 120.5 | – | – | N |
| AHam-070 | *Gmi* | 191914 | 1908 | 32,930,438 | 175,431 | 116.6 | – | – | N |
| AHam-071 | *Gmi* | 191914 | 1908 | 21,064,486 | 435,918 | 135.2 | – | – | N |
| AHam-072 | *Gmi* | 191914 | 1908 | 57,195,570 | 60,486 | 120.1 | – | – | N |
| AHam-073 | *Gmi* | 191914 | 1908 | 23,113,908 | 29,267 | 118.2 | – | – | N |
| AHam-074 | *Gmi* | 191914 | 1908 | 41,456,634 | 65,378 | 128.4 | – | – | N |
| AHam-075 | *Gmi* | 191914 | 1908 | 40,368,656 | 160,506 | 118.0 | – | – | N |
| AHam-076 | *Gmi* | 191914 | 1908 | 41,648,540 | 283,657 | 117.4 | – | – | N |
| AHam-077 | *Gmi* | 191914 | 1908 | 30,504,596 | 73,646 | 117.8 | – | – | N |
| AHam-078 | *Gmi** | 191914 | 1908 | 84,688,220 | 650,385 | 131.1 | 49.4 | 74 | N |
| AHam-079 | *Gmi** | 191914 | 1908 | 59,303,652 | 541,449 | 131.4 | 43.1 | 75 | N |
| AHam-080 | *Gmi* | 191914 | 1908 | 29,082,718 | 80,338 | 126.7 | – | – | N |
| AHam-081 | *Gmi* | 191914 | 1908 | 25,544,318 | 143,087 | 118.1 | – | – | N |
| AHam-082 | *Gmi* | 191914 | 1908 | 29,640,056 | 124,259 | 121.3 | – | – | N |
| AHam-083 | *Gmi** | 191914 | 1908 | 53,049,024 | 395,147 | 127.5 | 31.0 | 345 | N |
| AHam-084 | *Gmi* | 191914 | 1908 | 32,293,174 | 163,854 | 116.9 | – | – | N |

| | | | | | | | | | |
|---|---|---|---|---|---|---|---|---|---|
| AHam-086 | *Gmi* | 191914 | 1908 | 27,496,794 | 183,894 | 114.5 | – | – | N |
| AHam-087 | *Gmi* | 191914 | 1908 | 40,797,102 | 252,231 | 117.9 | – | – | N |
| AHam-088 | *Gmi* | 191914 | 1908 | 24,245,594 | 172,830 | 119.2 | – | – | N |
| AHam-089 | *Gmi** | 191914 | 1908 | 42,235,934 | 183,369 | 126.1 | 15.9 | 1,321 | N |
| AHam-090 | *Gmi* | 191914 | 1908 | 21,927,998 | 164,883 | 115.8 | – | – | N |
| AHam-091 | *Gmi* | 191914 | 1908 | 24,947,020 | 956,086 | 139.0 | – | – | N |
| AHam-092 | *Gmi* | 191914 | 1908 | 58,872,264 | 138,631 | 123.8 | – | – | N |
| AHam-093 | *Gmi* | 191914 | 1908 | 36,508,732 | 91,982 | 115.7 | – | – | N |
| AHam-094 | *Gmi* | 191914 | 1908 | 27,833,670 | 149,600 | 119.8 | – | – | N |
| AHam-095 | *Gmi* | 191914 | 1908 | 39,888,594 | 179,404 | 120.0 | – | – | N |
| AHam-096 | *Gmi* | 191914 | 1908 | 30,680,898 | 183,388 | 115.4 | – | – | N |
| AHam-001 | *Ela** | 191979 | 1908 | 43,865,858 | 5,233,960 | 110.9 | 123.3 | 3,203 | N |
| AHam-002 | *Ela** | 191979 | 1908 | 39,687,570 | 5,207,201 | 111.4 | 125.0 | 1,307 | N |
| AHam-003 | *Ela** | 191979 | 1908 | 47,617,604 | 6,897,211 | 107.2 | 160.3 | 493 | N |
| AHam-004 | *Ela** | 191979 | 1908 | 46,452,740 | 6,714,447 | 106.9 | 152.3 | 757 | N |
| AHam-005 | *Ela* | 191979 | 1908 | 44,303,360 | 6,800,917 | 106.7 | 149.8 | 934 | Y |
| AHam-006 | *Ela* | 191979 | 1908 | 49,330,014 | 7,889,896 | 102.6 | 169.1 | 510 | Y |
| AHam-007 | *Ela* | 191979 | 1908 | 36,993,366 | 4,784,150 | 106.5 | 106.4 | 3,709 | Y |
| AHam-008 | *Ela* | 191979 | 1908 | 14,065,872 | 1,014,171 | 99.9 | – | – | N |
| AHam-009 | *Ela* | 191979 | 1908 | 37,912,100 | 5,320,599 | 106.4 | 116.5 | 2,053 | Y |
| AHam-010 | *Ela* | 191979 | 1908 | 49,753,888 | 7,473,407 | 107.1 | 165.5 | 448 | Y |
| AHam-011 | *Ela** | 191979 | 1908 | 23,431,848 | 3,151,202 | 115.7 | 74.0 | 3,352 | N |
| AHam-012 | *Ela** | 191979 | 1908 | 30,140,562 | 4,106,775 | 116.5 | 101.3 | 991 | N |
| AHam-013 | *Ela* | 191979 | 1908 | 2,700,918 | 392,146 | 114.4 | – | – | N |
| AHam-014 | *Ela** | 191979 | 1908 | 47,645,886 | 6,742,663 | 107.7 | 148.9 | 715 | N |
| AHam-015 | *Ela** | 191979 | 1908 | 45,460,842 | 6,608,287 | 103.9 | 147.0 | 729 | N |
| AHam-016 | *Ela** | 191979 | 1908 | 11,434,556 | 1,619,409 | 119.2 | 44.5 | 5,891 | N |
| AHam-017 | *Ela** | 191979 | 1908 | 55,938,410 | 8,028,704 | 112.0 | 184.0 | 344 | N |
| AHam-018 | *Ela** | 191979 | 1908 | 47,495,738 | 6,513,658 | 107.5 | 140.1 | 900 | N |
| AHam-019 | *Ela* | 191979 | 1908 | 35,347,078 | 5,718,172 | 97.0 | 119.0 | 1,969 | Y |
| AHam-020 | *Ela* | 191979 | 1908 | 30,311,548 | 3,829,902 | 106.8 | 87.2 | 5,410 | Y |
| AHam-021 | *Ela* | 191979 | 1908 | 34,413,734 | 5,040,708 | 102.9 | 110.9 | 1,543 | Y |

| | | | | | | | | | |
|---|---|---|---|---|---|---|---|---|---|
| AHam-022 | *Ela* | 191979 | 1908 | 42,122,844 | 5,188,000 | 108.9 | 118.0 | 3,271 | Y |
| AHam-023 | *Ela** | 191979 | 1908 | 44,117,964 | 5,791,188 | 111.6 | 122.1 | 1,606 | N |
| AHam-024 | *Ela** | 191979 | 1908 | 36,895,112 | 4,459,887 | 115.2 | 104.5 | 1,459 | N |
| AHam-025 | *Ela** | 191979 | 1908 | 23,857,880 | 3,353,657 | 107.3 | 74.4 | 4,157 | N |
| AHam-026 | *Ela** | 191979 | 1908 | 26,575,210 | 4,022,360 | 105.4 | 96.6 | 2,032 | N |
| AHam-027 | *Ela** | 191979 | 1908 | 27,135,544 | 4,169,854 | 102.2 | 93.9 | 2,835 | N |
| AHam-028 | *Ela** | 191979 | 1908 | 16,241,648 | 2,838,051 | 104.9 | 72.9 | 3,361 | N |
| AHam-029 | *Ela** | 191979 | 1908 | 37,576,328 | 5,117,961 | 109.1 | 112.6 | 2,571 | N |
| AHam-030 | *Ela** | 191979 | 1908 | 42,411,444 | 5,371,136 | 106.6 | 112.4 | 2,420 | N |
| AHam-031 | *Ela* | 191979 | 1908 | 57,354,458 | 7,560,041 | 109.6 | 153.1 | 1,243 | Y |
| AHam-032 | *Ela* | 191979 | 1908 | 42,868,510 | 6,241,382 | 102.3 | 132.7 | 1,270 | Y |

**Table S2.** Sequencing statistics for all contemporary samples. Sample names are given in column one, species names in column two (*Gmi* = *G. minuta*, *Ela* = *E. laterofenestra*), and the year the sample was originally collected in column three. Column four lists the number of raw Illumina reads per sample. Column five gives the number of reads that mapped to the reference genome and passed filtering. Column seven lists the mean read length for mapped reads, while column eight gives the average read depth for each SNP used in most downstream analyses. Column nine gives the number of SNPs with missing genotypes for each individual. Column ten indicates whether or not the individual was used in the final analyses. An asterisk in the species column indicates the individual was identified as belonging to a cryptic species. Individuals involved in capture bait design are bolded. HH in parentheses in the included column indicates the sample was not included in the final analyses as its individual heterozygosity was three standard deviations above the species mean.

| Individual | Species | Year | Raw Reads | Mapped Reads | Mean Read Length | Mean SNP Depth | Missing SNPs | Included |
|---|---|---|---|---|---|---|---|---|
| CBat-001 | *Gmi* | 2018 | 5,902,132 | 237,744 | 146.1 | 9.81 | 2,848 | Y |
| CBat-002 | *Gmi* | 2018 | 9,088,734 | 461,740 | 145.4 | 20.96 | 415 | Y |
| CBat-003 | *Gmi* | 2018 | 8,020,142 | 356,330 | 146.2 | 14.01 | 1,335 | Y |
| CBat-004 | *Gmi** | 2018 | 16,152,122 | 283,524 | 142.0 | 14.81 | 2,077 | N |
| CBat-005 | *Gmi* | 2018 | 13,282,442 | 677,917 | 145.1 | 31.79 | 52 | Y |
| **CBat-006** | *Gmi* | 2018 | 4,387,168 | 175,174 | 145.4 | 11.22 | 1,790 | Y |
| **CBat-007** | *Gmi* | 2018 | 1,701,466 | – | – | – | – | N |
| CBat-008 | *Gmi** | 2018 | 29,134,322 | 466,964 | 145.4 | 42.62 | 167 | N |
| **CBat-009** | *Gmi* | 2018 | 5,903,038 | 195,941 | 145.3 | 14.41 | 1,036 | Y |
| CBat-010 | *Gmi* | 2018 | 31,621,308 | 675,834 | 145.1 | 18.06 | 671 | Y |
| CBat-011 | *Gmi* | 2018 | 7,343,204 | 290,749 | 145.9 | 13.21 | 1,521 | Y |
| CBat-012 | *Gmi* | 2018 | 9,774,714 | 511,168 | 145.1 | 20.89 | 383 | Y |
| CBat-013 | *Gmi* | 2018 | 16,319,760 | 674,327 | 143.7 | 40.46 | 18 | Y |
| CBat-014 | *Gmi* | 2018 | 19,330,092 | 730,057 | 144.9 | 59.52 | 7 | Y |
| CBat-015 | *Gmi* | 2018 | 11,879,844 | 526,129 | 142.7 | 41.38 | 16 | Y |
| CBat-016 | *Gmi* | 2018 | 23,231,700 | 920,959 | 145.2 | 64.22 | 8 | Y |
| CBat-017 | *Gmi** | 2018 | 18,562,608 | 342,588 | 145.5 | 27.49 | 362 | N |
| CBat-018 | *Gmi* | 2018 | 38,444,554 | 1,112,114 | 145.1 | 67.84 | 0 | Y |
| CBat-019 | *Gmi** | 2018 | 26,681,978 | 586,961 | 139.7 | 43.57 | 108 | N |
| CBat-020 | *Gmi* | 2018 | 18,622,874 | 594,719 | 143.0 | 37.73 | 26 | N (HH) |
| CBat-021 | *Gmi* | 2018 | 10,580,426 | 490,296 | 144.4 | 26.03 | 148 | Y |

| | | | | | | | | |
|---|---|---|---|---|---|---|---|---|
| CBat-022 | *Gmi* | 2018 | 8,516,576 | 381,088 | 143.8 | 24.63 | 224 | Y |
| CBat-023 | *Gmi* | 2018 | 10,118,370 | 223,476 | 145.4 | 8.63 | 3,193 | Y |
| CBat-024 | *Gmi* | 2018 | 10,791,794 | 528,227 | 145.6 | 23.94 | 274 | Y |
| **CBat-025** | *Gmi* | 2018 | 11,525,228 | 429,198 | 142.9 | 25.59 | 190 | Y |
| CBat-026 | *Gmi** | 2018 | 57,702,948 | 1,638,588 | 145.7 | 147.32 | 10 | N |
| CBat-027 | *Gmi* | 2018 | 17,955,856 | 639,214 | 143.6 | 37.85 | 30 | Y |
| CBat-028 | *Gmi* | 2018 | 12,061,584 | 367,589 | 143.6 | 20.28 | 675 | Y |
| CBat-029 | *Gmi** | 2018 | 31,024,962 | 922,956 | 145.8 | 83.73 | 20 | N |
| CBat-030 | *Gmi* | 2018 | 11,692,104 | 499,895 | 144.9 | 25.57 | 171 | Y |
| **CBat-031** | *Gmi** | 2018 | 22,385,248 | 617,085 | 141.5 | 48.79 | 134 | N |
| CBat-032 | *Gmi** | 2018 | 18,168,926 | 268,086 | 145.3 | 19.46 | 972 | N |
| **CBat-033** | *Gmi* | 2018 | 25,705,910 | 900,662 | 145.3 | 79.39 | 6 | Y |
| CBat-034 | *Gmi** | 2018 | 14,025,320 | 352,627 | 142.3 | 26.79 | 464 | N |
| CBat-035 | *Gmi* | 2018 | 22,187,598 | 1,040,879 | 144.3 | 46.95 | 17 | Y |
| CBat-036 | *Gmi* | 2018 | 7,827,090 | 379,156 | 145.1 | 21.28 | 326 | Y |
| **CBat-037** | *Gmi* | 2018 | 12,641,618 | 480,268 | 143.1 | 27.43 | 107 | Y |
| CBat-038 | *Gmi* | 2018 | 23,167,436 | 778,839 | 144.5 | 53.44 | 12 | Y |
| CBat-039 | *Gmi* | 2018 | 37,254,876 | 900,955 | 145.1 | 28.03 | 128 | Y |
| CBat-040 | *Gmi* | 2018 | 12,136,890 | 589,851 | 145.4 | 29.53 | 103 | Y |
| CBat-041 | *Gmi** | 2018 | 34,865,604 | 906,665 | 145.6 | 81.79 | 5 | N |
| CBat-042 | *Gmi* | 2018 | 10,889,974 | 365,845 | 143.3 | 25.66 | 146 | N (HH) |
| CBat-043 | *Gmi* | 2018 | 208,000,000 | 798,094 | 143.7 | 46.62 | 15 | Y |
| CBat-044 | *Gmi** | 2018 | 32,490,758 | 588,370 | 142.3 | 33.85 | 246 | N |
| CBat-045 | *Gmi* | 2018 | 30,672,824 | 862,301 | 143.7 | 48.68 | 28 | Y |
| CBat-046 | *Gmi* | 2018 | 35,371,982 | 923,280 | 144.2 | 36.92 | 22 | N (HH) |
| **CBat-047** | *Gmi** | 2018 | 22,415,756 | 603,941 | 142.0 | 45.20 | 157 | N |
| CBat-048 | *Gmi* | 2018 | 25,427,028 | 904,358 | 145.9 | 21.15 | 417 | Y |
| CBat-049 | *Gmi** | 2018 | 26,424,614 | 640,487 | 141.1 | 48.31 | 121 | N |
| **CBat-050** | *Gmi** | 2018 | 16,892,820 | 473,767 | 142.0 | 35.64 | 279 | N |
| CBat-051 | *Gmi* | 2018 | 17,653,876 | 675,288 | 142.9 | 44.72 | 4 | Y |
| **CBat-052** | *Gmi* | 2018 | 7,774,618 | 314,490 | 144.0 | 18.97 | 537 | Y |
| CBat-053 | *Gmi* | 2018 | 88,324,256 | 3,780,593 | 145.9 | 286.38 | 0 | Y |

| | | | | | | | | |
|---|---|---|---|---|---|---|---|---|
| CBat-054 | *Gmi** | 2018 | 16,850,160 | 593,278 | 145.6 | 56.30 | 26 | N |
| **CBat-056** | *Gmi* | 2018 | 12,965,714 | 507,744 | 144.4 | 30.97 | 100 | Y |
| **CBat-057** | *Gmi** | 2018 | 22,943,784 | 385,986 | 145.3 | 29.61 | 287 | N |
| CBat-058 | *Gmi* | 2018 | 263,242 | 1,380,938 | 145.7 | 25.08 | 157 | Y |
| **CBat-059** | *Gmi* | 2018 | 19,600,664 | 500,113 | 145.3 | 21.47 | 292 | Y |
| **CBat-060** | *Gmi* | 2018 | 49,481,200 | 1,289,515 | 144.3 | 45.79 | 2 | Y |
| CBat-061 | *Gmi* | 2018 | 10,468,402 | 457,363 | 144.7 | 25.97 | 142 | Y |
| CBat-062 | *Gmi* | 2018 | 8,952,554 | 456,011 | 145.0 | 24.61 | 187 | Y |
| CBat-063 | *Gmi* | 2018 | 35,139,292 | 919,926 | 145.0 | 36.20 | 25 | Y |
| CBat-064 | *Gmi** | 2018 | 5,989,132 | 247,632 | 142.1 | 22.50 | 737 | N |
| CBat-065 | *Gmi** | 2018 | 18,957,482 | 253,646 | 143.4 | 14.48 | 1,741 | N |
| CBat-066 | *Gmi* | 2018 | 115,000,000 | 861,136 | 143.4 | 50.16 | 4 | Y |
| CBat-067 | *Gmi* | 2018 | 22,317,544 | 812,511 | 143.3 | 50.82 | 3 | Y |
| CBat-068 | *Gmi* | 2018 | 1,913,772 | 76,167 | 144.8 | 8.02 | 3,354 | Y |
| **CBat-069** | *Gmi** | 2018 | 130,000,000 | 4,344,201 | 146.0 | 375.66 | 0 | N |
| CBat-070 | *Gmi* | 2018 | 14,841,780 | 708,284 | 144.1 | 43.21 | 16 | Y |
| **CBat-071** | *Gmi** | 2018 | 11,231,088 | 325,859 | 140.7 | 24.21 | 523 | N |
| CBat-072 | *Gmi* | 2018 | 9,918,372 | 216,094 | 143.7 | 12.45 | 2,157 | Y |
| CBat-073 | *Gmi** | 2018 | 25,668,310 | 562,980 | 145.6 | 39.57 | 163 | N |
| CBat-074 | *Gmi* | 2018 | 4,097,438 | 173,996 | 145.1 | 15.53 | 860 | Y |
| CBat-075 | *Gmi* | 2018 | 21,543,572 | 804,891 | 145.3 | 69.18 | 9 | Y |
| **CBat-076** | *Gmi* | 2018 | 7,363,236 | 291,662 | 144.4 | 20.15 | 349 | Y |
| CBat-077 | *Gmi* | 2018 | 89,304,452 | 3,368,522 | 145.6 | 270.59 | 0 | Y |
| **CBat-078** | *Gmi* | 2018 | 29,641,726 | 1,394,389 | 145.5 | 93.26 | 1 | Y |
| CBat-079 | *Gmi** | 2018 | 412,000,000 | 5,229,273 | 143.1 | 112.16 | 8 | N |
| CBat-080 | *Gmi* | 2018 | 14,960,598 | 725,421 | 144.3 | 41.47 | 26 | Y |
| CBat-081 | *Gmi* | 2018 | 35,286,414 | 1,015,408 | 144.6 | 62.27 | 2 | Y |
| CBat-082 | *Gmi* | 2018 | 14,101,848 | 632,019 | 143.5 | 40.48 | 27 | Y |
| CBat-083 | *Gmi* | 2018 | 5,580,308 | 271,615 | 144.4 | 20.55 | 455 | Y |
| CBat-084 | *Gmi** | 2018 | 27,751,598 | 588,497 | 142.4 | 37.74 | 206 | N |
| CBat-085 | *Gmi* | 2018 | 4,408,886 | 178,336 | 145.1 | 15.79 | 808 | Y |
| **CBat-086** | *Gmi* | 2018 | 42,321,424 | 1,508,994 | 145.3 | 127.60 | 3 | Y |

| | | | | | | | | |
|---|---|---|---|---|---|---|---|---|
| CBat-087 | *Gmi** | 2018 | 14,384,324 | 492,037 | 145.4 | 43.88 | 141 | N |
| **CBat-088** | *Gmi** | 2018 | 8,149,018 | 237,733 | 142.7 | 18.12 | 1,010 | N |
| CBat-089 | *Gmi* | 2018 | 43,946,156 | 1,751,293 | 144.8 | 125.04 | 0 | Y |
| **CBat-090** | *Gmi** | 2018 | 20,422,296 | 658,647 | 145.6 | 54.70 | 54 | N |
| **CBat-091** | *Gmi** | 2018 | 4,145,032 | 115,522 | 141.4 | 9.34 | 3,401 | N |
| CBat-092 | *Gmi* | 2018 | 26,702,316 | 1,104,594 | 145.6 | 97.82 | 0 | Y |
| CBat-093 | *Gmi* | 2018 | 8,804,152 | 411,051 | 144.7 | 24.06 | 201 | Y |
| CBat-094 | *Gmi* | 2018 | 15,755,210 | 715,397 | 143.9 | 38.16 | 11 | Y |
| CBat-095 | *Gmi* | 2018 | 7,441,928 | 395,153 | 144.7 | 17.79 | 565 | Y |
| **CBat-096** | *Gmi** | 2018 | 44,930,908 | 1,040,802 | 140.3 | 74.22 | 31 | N |
| CNas-001 | *Ela* | 2018 | 9,847,612 | 3,746,205 | 133.9 | 119.45 | 296 | Y |
| CNas-002 | *Ela* | 2018 | 7,777,716 | 2,825,200 | 133.8 | 83.53 | 661 | Y |
| CNas-003 | *Ela* | 2018 | 9,333,964 | 3,611,360 | 133.5 | 104.50 | 371 | Y |
| CNas-004 | *Ela* | 2018 | 12,399,778 | 4,188,297 | 135.3 | 117.50 | 336 | Y |
| **CNas-005** | *Ela* | 2018 | 6,035,646 | 2,289,381 | 135.0 | 73.98 | 1,135 | Y |
| CNas-006 | *Ela* | 2018 | 22,526,260 | 8,587,106 | 130.2 | 225.89 | 112 | Y |
| CNas-007 | *Ela* | 2018 | 8,381,580 | 3,115,711 | 132.2 | 93.46 | 533 | Y |
| CNas-008 | *Ela* | 2018 | 9,717,744 | 3,465,012 | 133.7 | 98.45 | 408 | Y |
| **CNas-009** | *Ela* | 2018 | 11,137,432 | 4,212,295 | 129.9 | 117.32 | 322 | Y |
| CNas-010 | *Ela* | 2018 | 8,391,106 | 3,050,212 | 132.5 | 89.37 | 581 | Y |
| **CNas-011** | *Ela* | 2018 | 7,640,134 | 2,849,410 | 134.4 | 89.89 | 577 | Y |
| CNas-012 | *Ela* | 2018 | 11,912,054 | 4,884,202 | 135.4 | 148.51 | 281 | Y |
| CNas-013 | *Ela* | 2018 | 7,287,498 | 2,838,217 | 132.0 | 86.67 | 710 | Y |
| CNas-014 | *Ela* | 2018 | 14,274,006 | 4,974,607 | 132.6 | 135.50 | 209 | Y |
| CNas-015 | *Ela* | 2018 | 9,167,122 | 3,373,565 | 129.8 | 94.14 | 335 | Y |
| CNas-016 | *Ela* | 2018 | 13,453,900 | 4,628,031 | 131.2 | 135.26 | 262 | Y |
| CNas-017 | *Ela* | 2018 | 23,418,658 | 8,237,783 | 124.0 | 188.74 | 74 | Y |
| CNas-018 | *Ela* | 2018 | 9,926,370 | 3,687,761 | 132.0 | 104.49 | 357 | Y |
| CNas-019 | *Ela* | 2018 | 6,852,358 | 2,608,170 | 133.2 | 81.70 | 848 | Y |
| CNas-020 | *Ela* | 2018 | 9,677,540 | 3,636,900 | 134.3 | 113.38 | 331 | Y |
| CNas-021 | *Ela* | 2018 | 17,940,428 | 6,507,196 | 132.7 | 175.41 | 146 | Y |
| CNas-022 | *Ela* | 2018 | 9,774,436 | 3,635,059 | 132.3 | 103.85 | 404 | Y |

| | | | | | | | | |
|---|---|---|---|---|---|---|---|---|
| CNas-023 | *Ela* | 2018 | 15,607,038 | 5,724,862 | 132.1 | 155.58 | 210 | Y |
| **CNas-024** | *Ela* | 2018 | 8,953,720 | 3,214,223 | 131.5 | 92.17 | 453 | Y |
| **CNas-025** | *Ela* | 2018 | 10,404,654 | 3,849,092 | 133.3 | 111.33 | 366 | Y |
| CNas-026 | *Ela* | 2018 | 3,280,328 | 1,397,134 | 131.8 | 44.45 | 2,829 | Y |
| CNas-027 | *Ela* | 2018 | 8,540,542 | 3,086,179 | 133.7 | 90.34 | 636 | Y |
| CNas-028 | *Ela* | 2018 | 11,081,680 | 3,745,539 | 132.9 | 102.74 | 325 | Y |
| **CNas-029** | *Ela* | 2018 | 6,568,992 | 2,282,851 | 132.9 | 65.35 | 961 | Y |
| CNas-030 | *Ela* | 2018 | 11,383,618 | 4,344,825 | 133.6 | 125.50 | 238 | Y |
| **CNas-031** | *Ela* | 2018 | 4,767,138 | 1,590,988 | 132.6 | 46.81 | 1,694 | Y |
| CNas-032 | *Ela* | 2018 | 9,324,836 | 3,238,380 | 132.5 | 88.56 | 449 | Y |
| **CNas-033** | *Ela* | 2018 | 24,826,742 | 9,647,288 | 132.9 | 273.06 | 45 | Y |
| **CNas-034** | *Ela* | 2018 | 8,409,906 | 3,054,493 | 134.6 | 96.19 | 386 | Y |
| CNas-035 | *Ela* | 2018 | 14,818,106 | 5,364,287 | 139.9 | 161.88 | 173 | Y |
| CNas-036 | *Ela* | 2018 | 10,793,360 | 3,940,301 | 132.9 | 119.93 | 260 | Y |
| CNas-037 | *Ela* | 2018 | 11,769,588 | 4,568,273 | 132.8 | 132.85 | 316 | Y |
| CNas-038 | *Ela* | 2018 | 15,653,980 | 6,211,083 | 133.1 | 180.46 | 133 | Y |
| CNas-039 | *Ela* | 2018 | 9,642,292 | 3,524,221 | 136.2 | 109.42 | 365 | Y |
| CNas-040 | *Ela* | 2018 | 13,621,270 | 5,013,671 | 127.4 | 135.05 | 236 | Y |
| CNas-041 | *Ela* | 2018 | 12,507,468 | 4,949,567 | 128.6 | 135.10 | 317 | Y |
| CNas-042 | *Ela* | 2018 | 18,517,068 | 7,246,427 | 131.3 | 203.44 | 109 | Y |
| **CNas-043** | *Ela** | 2018 | 3,019,600 | 1,055,301 | 130.9 | 29.90 | 0 | N |
| CNas-044 | *Ela* | 2018 | 10,297,836 | 3,826,389 | 131.0 | 105.28 | 409 | Y |
| CNas-045 | *Ela* | 2018 | 16,738,244 | 6,582,024 | 135.0 | 188.77 | 124 | Y |
| CNas-046 | *Ela* | 2018 | 9,601,584 | 3,421,109 | 134.8 | 103.61 | 385 | Y |
| **CNas-047** | *Ela* | 2018 | 8,072,980 | 2,775,957 | 132.1 | 78.25 | 619 | Y |
| CNas-048 | *Ela* | 2018 | 10,107,700 | 3,603,057 | 132.0 | 101.07 | 355 | Y |
| **CNas-049** | *Ela* | 2018 | 7,165,052 | 2,540,187 | 133.8 | 75.84 | 648 | Y |
| CNas-050 | *Ela* | 2018 | 10,114,962 | 3,543,049 | 132.6 | 101.89 | 387 | Y |
| CNas-051 | *Ela* | 2018 | 7,975,718 | 3,091,989 | 135.6 | 98.99 | 475 | Y |
| CNas-052 | *Ela* | 2018 | 3,996,198 | 1,654,108 | 131.9 | 49.27 | 2,072 | Y |
| **CNas-053** | *Ela* | 2018 | 10,139,166 | 3,762,679 | 132.9 | 116.28 | 358 | Y |
| **CNas-054** | *Ela* | 2018 | 14,780,096 | 5,372,574 | 130.1 | 140.53 | 152 | Y |

| | | | | | | | | |
|---|---|---|---|---|---|---|---|---|
| **CNas-055** | *Ela* | 2018 | 8,780,144 | 3,158,610 | 131.9 | 87.11 | 506 | Y |
| CNas-056 | *Ela* | 2018 | 20,407,972 | 8,080,687 | 135.7 | 230.99 | 113 | Y |
| CNas-057 | *Ela* | 2018 | 10,575,496 | 3,770,949 | 131.7 | 103.85 | 335 | Y |
| CNas-058 | *Ela* | 2018 | 10,802,510 | 3,713,073 | 132.1 | 106.80 | 264 | Y |
| CNas-059 | *Ela* | 2018 | 11,563,684 | 4,128,609 | 134.2 | 124.35 | 317 | Y |
| CNas-060 | *Ela* | 2018 | 7,756,000 | 2,827,386 | 132.8 | 85.81 | 539 | Y |
| **CNas-061** | *Ela* | 2018 | 5,239,690 | 2,008,118 | 134.2 | 64.08 | 1,459 | Y |
| **CNas-062** | *Ela* | 2018 | 7,175,666 | 2,801,142 | 133.2 | 83.21 | 872 | Y |
| CNas-063 | *Ela* | 2018 | 5,072,664 | 753,211 | 131.3 | – | – | N |
| CNas-064 | *Ela* | 2018 | 5,387,836 | 2,101,291 | 133.4 | 62.81 | 1,341 | Y |
| CNas-065 | *Ela* | 2018 | 10,442,838 | 3,889,012 | 133.9 | 116.74 | 321 | Y |
| CNas-066 | *Ela* | 2018 | 4,522,780 | 1,806,677 | 133.5 | 58.10 | 1,551 | Y |
| CNas-067 | *Ela* | 2018 | 9,730,012 | 3,363,591 | 130.9 | 90.45 | 465 | Y |
| CNas-068 | *Ela* | 2018 | 25,012,596 | 10,354,314 | 136.0 | 295.55 | 62 | Y |
| CNas-069 | *Ela* | 2018 | 7,491,468 | 2,573,840 | 133.3 | 71.75 | 592 | Y |
| CNas-070 | *Ela* | 2018 | 9,277,610 | 3,151,083 | 132.7 | 84.64 | 496 | Y |
| CNas-071 | *Ela* | 2018 | 9,034,794 | 3,314,920 | 134.2 | 104.01 | 415 | Y |
| **CNas-072** | *Ela* | 2018 | 10,126,052 | 3,719,649 | 134.1 | 115.38 | 270 | Y |
| CNas-073 | *Ela* | 2018 | 9,998,888 | 3,696,072 | 134.2 | 109.37 | 441 | Y |
| **CNas-074** | *Ela* | 2018 | 10,113,738 | 3,517,356 | 132.5 | 100.65 | 408 | Y |
| CNas-075 | *Ela* | 2018 | 10,996,264 | 3,632,754 | 132.5 | 101.63 | 356 | Y |
| CNas-076 | *Ela* | 2018 | 10,618,928 | 3,947,541 | 133.4 | 114.69 | 334 | Y |
| CNas-077 | *Ela* | 2018 | 7,844,078 | 2,868,955 | 133.2 | 81.32 | 693 | Y |
| CNas-078 | *Ela* | 2018 | 9,529,710 | 3,350,240 | 131.5 | 99.52 | 344 | Y |
| **CNas-079** | *Ela* | 2018 | 12,858,182 | 4,968,232 | 133.4 | 148.93 | 230 | Y |
| CNas-080 | *Ela* | 2018 | 8,833,818 | 3,268,515 | 134.8 | 99.35 | 526 | Y |
| CNas-081 | *Ela* | 2018 | 14,388,588 | 4,902,317 | 132.9 | 139.37 | 205 | Y |
| CNas-082 | *Ela* | 2018 | 9,232,088 | 3,215,528 | 132.9 | 94.02 | 392 | Y |
| CNas-083 | *Ela* | 2018 | 13,314,462 | 4,749,304 | 134.0 | 135.50 | 210 | Y |
| **CNas-084** | *Ela* | 2018 | 12,026,774 | 4,424,460 | 133.9 | 128.99 | 308 | Y |
| CNas-085 | *Ela* | 2018 | 10,952,816 | 4,025,567 | 132.5 | 120.94 | 306 | Y |
| CNas-086 | *Ela* | 2018 | 8,718,996 | 3,135,326 | 133.3 | 89.73 | 515 | Y |

| | | | | | | | | |
|---|---|---|---|---|---|---|---|---|
| CNas-087 | *Ela* | 2018 | 9,940,868 | 3,312,072 | 133.5 | 95.01 | 384 | Y |
| CNas-088 | *Ela* | 2018 | 9,644,796 | 3,687,114 | 135.4 | 114.23 | 337 | Y |
| CNas-089 | *Ela* | 2018 | 9,017,988 | 3,215,113 | 134.5 | 94.95 | 478 | Y |
| **CNas-090** | *Ela* | 2018 | 9,115,088 | 3,371,165 | 135.0 | 107.16 | 453 | Y |
| **CNas-091** | *Ela* | 2018 | 7,934,268 | 2,943,118 | 134.7 | 92.89 | 654 | Y |
| CNas-092 | *Ela* | 2018 | 21,142,420 | 8,326,670 | 134.8 | 239.86 | 80 | Y |
| CNas-093 | *Ela* | 2018 | 5,990,272 | 2,518,930 | 128.9 | 70.74 | 1,170 | Y |
| **CNas-094** | *Ela* | 2018 | 9,442,510 | 3,409,750 | 133.6 | 99.73 | 393 | N (HH) |
| CNas-095 | *Ela* | 2018 | 20,182,258 | 7,892,929 | 136.4 | 248.13 | 48 | N (HH) |
| CNas-096 | *Ela* | 2018 | 15,548,260 | 5,854,144 | 130.8 | 168.72 | 188 | Y |

**Table S3.** Demographic modeling results for each of the 15 demographic models fit to the contemporary SFS for *G. minuta*. The number of parameters and log likelihood are reported for each model, along with ΔAIC compared to the top model (model AIC - top AIC). For the top model, AIC is also reported in parentheses. The row with the top model is highlighted in gray.

| Model | Number of Parameters | Log Likelihood | ΔAIC |
|---|---|---|---|
| Constant $N_e$ | 1 | -18,372.83 | 2,805.24 |
| Recent abrupt change in $N_e$ | 3 | -18,366.26 | 2,796.10 |
| Recent gradual change in $N_e$ | 3 | -18,366.81 | 2,797.20 |
| Historical abrupt change in $N_e$ | 3 | -18,341.88 | 2,747.34 |
| Historical gradual change in $N_e$ | 3 | -18,344.38 | 2,752.34 |
| Ancient abrupt change in $N_e$ | 3 | -17,125.21 | 314.00 |
| Ancient gradual change in $N_e$ | 3 | -17,213.51 | 490.60 |
| Recent and historical abrupt changes in $N_e$ | 5 | -18,341.88 | 2,751.34 |
| Recent and historical gradual changes in $N_e$ | 5 | -18,343.87 | 2,755.32 |
| Recent and ancient abrupt changes in $N_e$ | 5 | -16,968.06 | 3.70 |
| Recent and ancient gradual changes in $N_e$ | 5 | -16,966.21 | 0 (33,942.42) |
| Historical and ancient abrupt changes in $N_e$ | 5 | -16,968.24 | 4.06 |
| Historical and ancient gradual changes in $N_e$ | 5 | -16,966.25 | 0.08 |
| Recent, historical, and ancient abrupt changes in $N_e$ | 7 | -16,967.99 | 7.56 |
| Recent, historical, and ancient gradual changes in $N_e$ | 7 | -16,966.24 | 4.06 |

**Table S4.** Demographic modeling results for each of the 15 demographic models fit to the contemporary SFS for *E. laterofenestra*. The number of parameters and log likelihood are reported for each model, along with ΔAIC compared to the top model (model AIC - top AIC). For the top model, AIC is also reported in parentheses. The row with the top model is highlighted in gray.

| Model | Number of Parameters | Log Likelihood | ΔAIC |
|---|---|---|---|
| Constant $N_e$ | 1 | -164,614.52 | 9,230.38 |
| Recent abrupt change in $N_e$ | 3 | -163,853.90 | 7,713.14 |
| Recent gradual change in $N_e$ | 3 | -163,852.39 | 7,710.12 |
| Historical abrupt change in $N_e$ | 3 | -163,856.03 | 7,717.40 |
| Historical gradual change in $N_e$ | 3 | -163,852.62 | 7,740.58 |
| Ancient abrupt change in $N_e$ | 3 | -164,146.99 | 8,299.32 |
| Ancient gradual change in $N_e$ | 3 | -164,022.29 | 8,049.92 |
| Recent and historical abrupt changes in $N_e$ | 5 | -163,851.48 | 7,712.30 |
| Recent and historical gradual changes in $N_e$ | 5 | -163,851.56 | 7,712.46 |
| Recent and ancient abrupt changes in $N_e$ | 5 | -159,998.85 | 7.04 |
| Recent and ancient gradual changes in $N_e$ | 5 | -159,995.33 | 0 (320,000.66) |
| Historical and ancient abrupt changes in $N_e$ | 5 | -160,000.31 | 9.96 |
| Historical and ancient gradual changes in $N_e$ | 5 | -159,998.43 | 6.20 |
| Recent, historical, and ancient abrupt changes in $N_e$ | 7 | -159,999.29 | 11.92 |
| Recent, historical, and ancient gradual changes in $N_e$ | 7 | -159,995.73 | 4.80 |

**Table S5.** $N_e$ estimates for both species, generated from demographic modeling with momi2. Results from the most parsimonious model fit to the historical SFS. Estimates of ancient $N_e$, historical $N_e$, ($N_e$ of the historical population), and the timing of the ancient and historical population size change (in years before present). Bootstrapped 95% CIs are provided in brackets.

| | **Ancient $N_e$** | **Historical $N_e$** | **Time of Ancient $N_e$ Change** | **Time of Historical $N_e$ Change** |
|---|---|---|---|---|
| ***G. minuta*** | 71,209 [65,182 - 75,454] | 86 [8 - 7,274,347,000] | 99,997 [99,465 - 100,000] | 120 [110 - 483] |
| ***E. laterofenestra*** | 804,749 [787,924 - 824,803] | 10,000,000,000 [1,515,974,166 - 10,000,000,000] | 99,999 [98,972 - 100,000] | — |

**Table S6.** $N_e$ estimates for both species, generated from demographic modeling with momi2. Results from the most parsimonious model fit to the historical and contemporary SFS. Estimates of ancient $N_e$, historical $N_e$ ($N_e$ of the historical population), contemporary $N_e$, and the timing of the ancient, historical, and recent population size change (in years before present). Bootstrapped 95% CIs are provided in brackets.

| | **Ancient $N_e$** | **Historical $N_e$** | **Contemporary $N_e$** | **Time of Ancient $N_e$ Change** | **Time of Historical $N_e$ Change** | **Time of Recent $N_e$ Change** |
|---|---|---|---|---|---|---|
| ***G. minuta*** | 67,528 [43,012 - 74,798] | 5,083 [2,339 - 12,040] | 10,000,000,000 [245,758 - 10,000,000,000] | 153,112 [139,983 - 287,003] | 499 [199 - 500] | 100 [44 - 100] |
| ***E. laterofenestra*** | 688,430 [610,207 - 707,119] | 2,258 [1,032 - 715,572] | 7,455 [1,303 - 16,933,049] | 633,522 [600,151 - 1,000,000] | 339 [186 - 494] | 79 [7 - 100] |

**Table S7.** Demographic modeling results for each of the 7 demographic models fit to the historical SFS for *G. minuta*. The number of parameters and log likelihood are reported for each model, along with ΔAIC compared to the top model (model AIC - top AIC). For the top model, AIC is also reported in parentheses. The row with the top model is highlighted in gray.

| **Model** | **Number of Parameters** | **Log Likelihood** | **ΔAIC** |
|---|---|---|---|
| Constant $N_e$ | 1 | -9,636.23 | 1,317.92 |
| Historical abrupt change in $N_e$ | 3 | -9,620.49 | 1,290.44 |
| Historical gradual change in $N_e$ | 3 | -9,622.10 | 1,293.66 |
| Ancient abrupt change in $N_e$ | 3 | -8,999.29 | 48.04 |
| Ancient gradual change in $N_e$ | 3 | -9,065.52 | 180.50 |
| Historical and ancient abrupt changes in $N_e$ | 5 | -8,973.27 | 0 (17,956.54) |
| Historical and ancient gradual changes in $N_e$ | 5 | -8,979.99 | 13.44 |

**Table S8.** Demographic modeling results for each of the 7 demographic models fit to the historical SFS for *E. laterofenestra*. The number of parameters and log likelihood are reported for each model, along with ΔAIC compared to the top model (model AIC - top AIC). For the top model, AIC is also reported in parentheses. The row with the top model is highlighted in gray.

| Model | Number of Parameters | Log Likelihood | ΔAIC |
|---|---|---|---|
| Constant $N_e$ | 1 | -49,024.41 | 1,762.40 |
| Historical abrupt change in $N_e$ | 3 | -49,021.11 | 1,759.80 |
| Historical gradual change in $N_e$ | 3 | -49,019.85 | 1,757.28 |
| Ancient abrupt change in $N_e$ | 3 | -48,141.21 | 0 (96,288.42) |
| Ancient gradual change in $N_e$ | 3 | -48,196.79 | 111.16 |
| Historical and ancient abrupt changes in $N_e$ | 5 | -48,141.20 | 3.98 |
| Historical and ancient gradual changes in $N_e$ | 5 | -48,196.61 | 114.80 |

**Table S9.** Demographic modeling results for each of the 15 demographic models fit to the historical and contemporary SFS for *G. minuta*. The number of parameters and log likelihood are reported for each model, along with ΔAIC compared to the top model (model AIC - top AIC). For the top model, AIC is also reported in parentheses. The row with the top model is highlighted in gray.

| Model | Number of Parameters | Log Likelihood | ΔAIC |
|---|---|---|---|
| Constant $N_e$ | 1 | -31,120.67 | 2,869.44 |
| Recent abrupt change in $N_e$ | 3 | -31,120.60 | 2,873.30 |
| Recent gradual change in $N_e$ | 3 | -31,120.61 | 2,873.32 |
| Historical abrupt change in $N_e$ | 3 | -31,112.63 | 2,857.36 |
| Historical gradual change in $N_e$ | 3 | -31,114.78 | 2,861.66 |
| Ancient abrupt change in $N_e$ | 3 | -29,950.54 | 533.18 |
| Ancient gradual change in $N_e$ | 3 | -30,068.59 | 769.28 |
| Recent and historical abrupt changes in $N_e$ | 5 | -31,109.69 | 2,855.48 |
| Recent and historical gradual changes in $N_e$ | 5 | -31,100.99 | 2,838.08 |
| Recent and ancient abrupt changes in $N_e$ | 5 | -29,899.13 | 434.36 |
| Recent and ancient gradual changes in $N_e$ | 5 | -29,993.50 | 623.10 |
| Historical and ancient abrupt changes in $N_e$ | 5 | -29,725.59 | 87.28 |
| Historical and ancient gradual changes in $N_e$ | 5 | -29,833.61 | 303.32 |
| Recent, historical, and ancient abrupt changes in $N_e$ | 7 | -29,679.95 | 0 (59,373.90) |
| Recent, historical, and ancient gradual changes in $N_e$ | 7 | -29,834.92 | 309.94 |

**Table S10.** Demographic modeling results for each of the 15 demographic models fit to the historical and contemporary SFS for *E. laterofenestra*. The number of parameters and log likelihood are reported for each model, along with ΔAIC compared to the top model (model AIC - top AIC). For the top model, AIC is also reported in parentheses. The row with the top model is highlighted in gray.

| Model | Number of Parameters | Log Likelihood | ΔAIC |
|---|---|---|---|
| Constant $N_e$ | 1 | -232,942.24 | 12,337.38 |
| Recent abrupt change in $N_e$ | 3 | -231,053.36 | 8,563.62 |
| Recent gradual change in $N_e$ | 3 | -231,050.92 | 8,558.74 |
| Historical abrupt change in $N_e$ | 3 | -231,055.30 | 8,567.50 |
| Historical gradual change in $N_e$ | 3 | -231,052.16 | 8,561.22 |
| Ancient abrupt change in $N_e$ | 3 | -231,371.74 | 9,200.38 |
| Ancient gradual change in $N_e$ | 3 | -232,189.55 | 10,836.00 |
| Recent and historical abrupt changes in $N_e$ | 5 | -231,052.84 | 8,566.58 |
| Recent and historical gradual changes in $N_e$ | 5 | -231,049.40 | 8,559.70 |
| Recent and ancient abrupt changes in $N_e$ | 5 | -227,767.60 | 1,996.10 |
| Recent and ancient gradual changes in $N_e$ | 5 | -231,049.40 | 8,559.70 |
| Historical and ancient abrupt changes in $N_e$ | 5 | -230,540.68 | 7,542.26 |
| Historical and ancient gradual changes in $N_e$ | 5 | -231,940.73 | 10,342.36 |
| Recent, historical, and ancient abrupt changes in $N_e$ | 7 | -226,767.55 | 0 (453,549.10) |
| Recent, historical, and ancient gradual changes in $N_e$ | 7 | -228,094.44 | 2,653.78 |

**SI References:**


1. O. A. Ali et al., RAD Capture (Rapture): Flexible and efficient sequence-based genotyping. *Genet.* 202, 389-400 (2016).
2. R. L. Roberts, "Whole genome sequencing century-old ethanol-preserved Philippine fishes," thesis, Texas A&M University - Corpus Christi (2023); https://hdl.handle.net/1969.6/97716.
3. J. Catchen, P. A. Hohenlohe, S. Bassham, A. Amores, W. A. Cresko, Stacks: An analysis tool set for population genomics. *Mol. Ecol.* 22, 3124-3140 (2013).
4. C. E. Bird, dDocentHPC: A pipeline to trim, assemble, map, and genotype reduced representation genomic data (2022); https://github.com/cbirdlab/dDocentHPC.
5. J. B. Puritz, C. M. Hollenbeck, J. R. Gold, *dDocent*: A RADseq, variant-calling pipeline designed for population genomics of non-model organisms. *PeerJ* 2, e431 (2014).
6. A. M. Bolger, M. Lohse, B. Usadel, Trimmomatic: A flexible trimmer for Illumina sequence data. *Bioinformatics* 30, 2114-2020 (2014).
7. Z. Chong, J. Ruan, C. I. Wu, Rainbow: An integrated tool for efficient clustering and assembling RAD-seq reads. *Bioinformatics* 28, 2732-2737 (2012).
8. M. Vasimuddin, S. Misra, H. Li, S. Aluru in *IEEE International Parallel and Distributed Processing Symposium (IPDPS)* 314-324 (2019).
9. P. Danecek et al., Twelve years of SAMtools and BCFtools. *GigaScience* 10, giab008 (2021).
10. E. Garrison, G. Marth, Haplotype-based variant detection from short-read sequencing. arXiv:1207.3907 [genomics] (2012).
11. P. Danecek et al., The variant call format and VCFtools. *Bioinformatics* 27, 2156-2158 (2011).
12. S. C. Willis, C. M. Hollenbeck, J. B. Puritz, J. R. Gold, D. S. Portnoy, Haplotyping RAD loci: An efficient method to filter paralogs and account for physical linkage. *Mol. Ecol. Resour.* 17, 955-965 (2017).
13. A. Gnirke et al., Solution hybrid selection with ultra-long oligonucleotides for massively parallel targeted sequencing. *Nat. Biotechnol.* 27, 182-189 (2009).
14. S. Andrews, FastQC: A quality control tool for high throughput sequence data (2010); https://www.bioinformatics.barbraham.ac.uk/projects/fastqc/.
15. S. Chen, Y. Zhou, Y. Chen, J. Gu, fastp: An ultra-fast all-in-one FASTQ preprocessor. *Bioinformatics* 34, i884-i890 (2018).
16. B. Bushnell, BBTools (2019); https://sourceforge.net/projects/bbmap/.
17. S. W. Wingett, S. Andrew, FastQ Screen: A tool for multi-genome mapping and quality control. *F1000Research* 7, 1338 (2018).
18. B. Langmead, S. L. Salzburg, Fast gapped-read alignment with Bowtie 2. *Nat. Methods* 9, 357-359 (2012).
19. S. Purcell et al., PLINK: A tool set for whole-genome association and population-based linkage analyses. *Am. J. Hum. Genet.* 81, 559-575 (2007).
20. Sparks, J. S. & Chakrabarty, P. A new species of ponyfish (Teleostei: Leognathidae: *Photoplagios*) from the Philippines. *Copeia* 3, 622-629 (2007).
21. E. L. Petrou et al., Intraspecific DNA contamination distorts subtle population structure in a marine fish: Decontamination of herring samples before restriction site-associated

sequencing and its effects on population genetic statistics. *Mol. Ecol. Resour.* 19, 1131-1143 (2019).
22. H. Jónsson, A. Ginolhac, M. Schubert, P. L. F. Johnson, L. Orlando, mapDamage2.0: Fast approximate Bayesian estimates of ancient DNA damage parameters. *Bioinformatics* 29, 1682-1684 (2013).
23. S. Dolenz et al., Unravelling reference bias in ancient DNA datasets. *Bioinformatics* 40, btae436 (2024).
24. T. Günther, C. Nettelblad, The presence and impact of reference bias on population genomic studies of prehistoric human populations. *PLoS Genet.* 15, e1008302 (2019).
25. M. Kardos, R. S. Waples, Low-coverage sequencing and Wahlund effect severely bias estimates of inbreeding, heterozygosity and effective population size in North American wolves. *Mol. Ecol.*, e17415 (2024).
26. T. Maruki, M. Lynch, Genotype calling from population-genomic sequencing data. *G3: Genes Genomes Genet.* 7, 1393-1404 (2017).
27. J. T. Thorson et al., Identifying direct and indirect associations among traits by merging phylogenetic comparative methods and structural equation models. *Methods Ecol. Evol.* 14, 1259-1275 (2023).
28. M. Roesti, D. Moser, D. Berner, Recombination in the threespine stickleback genome - patterns and consequences. *Mol. Ecol.* 22, 3014-3027 (2013).
29. C. Feng et al., Moderate nucleotide diversity in the Atlantic herring is associated with a low mutation rate. *eLife* 30, e23907 (2017).
30. J. Kamm, J. Terhost, R. Durbin, Y. S. Song, Efficiently inferring the demographic history of many populations with allele count data. *J. Amer. Statist. Assoc.* 115, 1472-1487 (2020).
